# Liraglutide treatment reverses unconventional cellular defects in induced pluripotent stem cell-derived β cells harboring a partially functional WFS1 variant

**DOI:** 10.1101/2024.08.06.606801

**Authors:** Silvia Torchio, Gabriel Siracusano, Federica Cuozzo, Valentina Zamarian, Silvia Pellegrini, Fabio Manenti, Riccardo Bonfanti, Giulio Frontino, Valeria Sordi, Raniero Chimienti, Lorenzo Piemonti

## Abstract

**Aims/hypothesis:** Wolfram Syndrome 1 (WS1) is a rare genetic disorder characterized by very heterogeneous clinical manifestations caused by variants of the *WFS1* gene, which encodes for the Endoplasmic Reticulum (ER) protein Wolframin, involved in cellular stress response, Ca^2+^ handling and autophagy. Given the central role of Wolframin, elucidating the impact of *WFS1* variants on cell functions is crucial to provide an association with clinical phenotypes. Therefore, as the understanding of patient-specific defects may also help to develop targeted therapeutic approaches, here we aimed at elucidating the impact on β cell function of the c.316-1G>A mutation harboring the partially functional Wolframin that we have previously characterized, and the molecular changes following treatment with the glucagon-like peptide 1 receptor (GLP-1R) agonist liraglutide.

**Methods:** We previously generated patient-derived iPSCs (WFS1) and isogenic line in which the c.316-1G>A mutation was genetically corrected (WFS1^wt/757A>T^), thus performed molecular analysis, including single cell RNAseq (scRNAseq), and functional studies on iPSC-derived β cell (iBeta). Calcium flux imaging and dynamic perifusion assays were used to test glucose responsiveness of iBeta. Treatment with liraglutide was performed to investigate effects on glucose-stimulated insulin secretion (GSIS), unfolded protein response (UPR), autophagy and apoptosis.

**Results:** We found that both WFS1 and WFS1^wt/757A>T^ iBeta efficiently differentiated *in vitro* into pancreatic lineage, but WFS1 showed less mature endocrine phenotype, reduced glucose responsiveness and impaired insulin secretion compared to WFS1^wt/757A>T^ counterpart. The Ca^2+^ dynamics were altered in WFS1 iBeta as Ca^2+^ oscillations after glucose challenge were not synchronized mainly due to the *CACNA1D* and *SNAP25* downmodulation. Reduced insulin secretion was correlated with a decrease in PC1/3 levels and overall increase of *RGS4* expression in WFS1 iBeta, whereas secretory defects correlated with accelerated autophagic flux. While the functional residual Wolframin in WFS1 iBeta controlled short-term ER stress, prolonged insults or inflammation highlighted ineffective UPR that make the cells unable to escape apoptosis. Interestingly, treatment with liraglutide restored the Ca^2+^ fluxes and secretory impairments, increasing glucose responsiveness and insulin release of WFS1 iBeta, while protecting these cells from cellular stress and inflammation-induced apoptosis.

**Conclusion/interpretation:** Our data highlighted alterations of key cellular pathways involved in WS1 β cell maturation and GSIS and how the pharmacological targeting of the GLP-1/GLP-1R axis was able to restore the physiologic phenotype. This study points out the need to understand the patient-specific molecular determinants associated with *WFS1* variants, to design effective therapies to treat the disease.

## Introduction

Wolfram Syndrome 1 (WS1) is a rare autosomal recessive, multisystemic disease, with a very low estimated prevalence of around 1:700000 in the general population. The onset typically manifests in the first and second decades of life with diabetes mellitus (DM) and optic atrophy (OA). As the disease progresses, additional complications emerge, including hearing loss, diabetes insipidus, urinary dysfunctions, and ultimately leading to premature death due to central respiratory failure [1–3]. WS1 clinical manifestations are caused by pathogenic variants of the *WFS1* gene, which encodes for a transmembrane endoplasmic reticulum (ER) protein known as Wolframin. In pancreatic cells, Wolframin preserves normal β cell physiology by promoting insulin biosynthesis and negatively regulating ER stress-inducible pro-apoptotic factors [4]. ER stress is regulated by three unfolded protein response (UPR) signaling pathways mediated by three ER membrane sensors: protein kinase RNA-like endoplasmic reticulum kinase (PERK), inositol-requiring enzyme 1 (IRE1α), and the activating transcription factor 6 (ATF6). These pathways coordinate homeostasis recovery in stressed cells and mediate the switch between survival and apoptosis [5]. Misregulation of ER stress response in WS1-affected cells was reported as a hallmark of the disorder, as Wolframin deficiency upregulated the three UPR pathways [6]. Wolframin exerts a relevant role in Ca^2+^ homeostasis because it could function as or activate an ER Ca^2+^ channel [7] and promotes Ca^2+^ transfer between the ER and mitochondria [8]. Ca^2+^ signaling is of utmost importance in β cells, where inter-organelle communication and insulin secretion are regulated by precise ion transfer in and out of the cytoplasm, which is directly and indirectly handled by Wolframin [9, 10]. It has been reported that the loss of WSF1 and the consequent alteration of the Ca^2+^ dynamics activated autophagy and mitophagy to supply the mitochondrial metabolism and increase energy production [11, 12]. WS1 β cells also display secondary dysfunctions such as diminished insulin secretory response to glucose, elevated proinsulin/insulin ratio, reduced islet number *in vivo* and poor survival upon exogenous stress application [13–15].

Given the central role of Wolframin in several cellular processes, elucidating the impact of *WFS1* variants on cell function is crucial to provide an association with the clinical phenotype. Indeed, there are no preventive or curative options for WS1, but several drug repurposing–based approaches are currently under investigation to manage the clinical manifestations, including chemical chaperones, ER calcium stabilizers, ER stress targeting [1, 4, 16]. Although these drugs showed promising results in both preclinical and clinical settings, little is known about the exact mechanisms by which WS1 β cells degenerate, and the paucity of information underlies a severe constraint on putative pharmacological targets.

Increasing evidence suggests Glucagon-like peptide 1 receptor (GLP-1R) agonists as alternative therapeutic options for their pleiotropic effects on ER stress, autophagy, Ca^2+^ handling and insulin secretion [17]. An off-label clinical trial employing liraglutide showed that it is well tolerated by pediatric patients and potentially slows down the progressive degeneration, both at the pancreatic endocrine and neuronal levels [18]. Accordingly, long-lasting GLP-1R agonist dulaglutide reversed impaired glucose tolerance in Wolframin-deficient mice, and exenatide and dulaglutide improved β cell function, preventing apoptosis in different human Wolframin-deficient models including inducing Pluripotent Stem Cell (iPSC)-derived β cells from WS1 patients [19].

We have previously characterized a WS1 patient carrying two pathogenic compound heterozygous mutations in *WFS1* gene: c.316-1G>A, leading to splice acceptor site disruption upstream exon 4, and c.757A>T, introducing a premature stop codon in exon 7 [18, 20, 21]. In this patient, the allele carrying the c.316-1G>A mutation originates two premature termination codon (PTC)-containing alternative splicing transcripts (c.316del and c.316-460del), that are regulated by nonsense mediated decay (NMD), and two open reading frame (ORF)-conserving mRNAs (c.271-513del and c.316-456del) leading to N-terminally truncated polypeptides, that retain the functional C-terminal domain, partially preserving the ER-stress response in iPSC-derived iBeta. We found that exogenous stress induced the accumulation of PTC-carrying mRNAs, leading to cell death [21]. The patient is currently under liraglutide treatment and showed promising improvement of clinical symptoms [18]. In this work, we aimed at analyzing the functional impact of the characterized mutations by exploring the potential cellular mechanisms involved in ꞵ cell dysfunction by using WS1 patient iPSC-derived ꞵ cell. Moreover, we attempted to elucidate whether these mechanisms might be targeted by liraglutide treatment, thus explaining the therapeutic efficacy of this drug.

## Methods and Materials

### Cell culture

The two WFS1 (#42 and #43) and three WFS1^wt/757A>T^ (#B10, #C6 and #F4) iPSCs clones used in this study were previously described in [21]. All the iPSC lines were routinely tested for mycoplasma contamination by using MycoAlert™ Mycoplasma Detection Kit by Lonza, according to the manufacturer’s instructions. Differentiation into pancreatic β cells was carried out in adhesion according to the published *in vitro* protocol [22]. At the end of differentiation, iPSC-derived β cells (iBeta) were detached using 0.5 mM EDTA and transferred into 60 mm petri dishes for re-clustering into pseudo-islets on an orbital shaker at 55 rpm. Pseudo-islets were maintained in suspension culture until their use, into CMRL 1066 (Mediatech) supplemented with 10% fetal bovine serum (FBS) (Lonza), 1% Penicillin/Streptomycin (Pen/Strep), 1% L-Glutamine (Euroclone), 10 μM Alk5i II, 1 μM T3, 10 mM Nicotinamide (Sigma), 10 μM H1152 (Euroclone), 100 U/ml DNase I (Sigma). To study the impact of the exogenous stress, iBeta were treated with 50 or 500 nM Thapsigargin for 8h or 16h, or with 1000U/ml IFNγ, 10 ng/ml TNFɑ, 50 U/ml IL-1β for 48h. To study autophagy, iBeta were treated with 100 nM bafilomycin for 24h. For the apoptosis studies, 1 μM liraglutide was added to the medium a day before and throughout the time of either TG or pro-inflammatory cytokine treatment.

### Flow cytometry

Pseudo-islets were dissociated with Trypsin, reduced to a single cell suspension and stained with Live/Dead Fixable Violet stain kit (ThermoScientifics) to exclude dead cells. Cells were fixed by using Cytofix/Cytoperm (BD Biosciences), then permeabilized with Phosflow™ Perm Buffer III (BD Biosciences) for intracellular staining with conjugated antibodies for 30’ at 4°C. A list of all antibodies used for FACS staining can be found in **ESM Table 1**. For apoptosis assay, P.I. (Sigma) and FITC Annexin V Apoptosis Detection Kit I (BD Biosciences) were used according to the manufacturer’s instructions. All flow cytometry analyses were performed on a FACS Canto (BD Biosciences) and results were analyzed with FlowJo™ Software V.10.

### Immunofluorescence

The iBeta were fixed with 4% paraformaldehyde in PBS for 20’ at room temperature (RT), then treated for de-masking with 15 mM glycine for 5’ at RT. After blocking/permeabilization with PBS supplemented with 0.4% Triton X-100, 2% BSA, 5% FBS for 45’ at RT, iBeta were incubated overnight at 4°C with the appropriate primary antibody (**ESM Table 1**), following incubation with secondary antibodies (**ESM Table 1**) for 1h at RT. Primary and secondary antibodies were diluted in PBS supplemented with 2% BSA. Nuclei were counterstained with Hoechst 33342. Images were acquired using the Olympus FluoVIEW FV 3000 (VivaScope Research) confocal microscope and analyzed using the Fiji software v.1.52p.

### RNA extraction, retrotranscription and RT-qPCR

The total RNA was extracted by the mirVana Isolation Kit (Ambion), following the treatment with TURBO^TM^ DNAse (Thermo Fisher) according to manufacturers’ instructions. About 1µg of RNA was reverse transcribed by using the SuperScript IV RT (Invitrogen). Predesigned TaqMan Array microfluidic cards (Applied Biosystems) with a total of 48 endocrine/β cell markers (**ESM Table 2**) were used for differential gene expression analysis on 4 preparations of WFS1 and WFS1^wt/757A>T^ pancreatic progenitors (day 14), and 7-8 preparations of WFS1 and WFS1^wt/757A>T^ iBeta (day 24). The gene-specific forward/reverse primers used for 96-well RT-qPCR experiments with PowerUp Green Master Mix (Applied Biosystems) are listed in **ESM Table 3**. The TaqMan array card and SYBR-based 96-well RT-qPCRs were performed on QuantStudio 12K Flex (Applied Biosystems) platform, according to the manufacturer’s instructions, setting up the amplification protocol with the annealing temperature of 60°C. Results are reported as normalized over the expression of the housekeeping gene *GAPDH* and displayed as absolute normalized quantity (2^-ΔCt^) or as a fold over a control (2^-ΔΔCt^).

### Single cell RNA sequencing

The WFS1 and WFS1^wt/757A>T^ iBeta from two clones/line were disaggregated by trypsin (Lonza) and captured using the droplet-based Chromium 10X platform, kit version 3. Single cell RNA sequencing (scRNAseq) was performed on Illumina NovaSeq, obtaining about 150 millions of paired end reads per sample. Cellular barcodes corresponding to good-quality cells were identified and extracted using the UMItool pipeline, then reads were aligned on the human genome hg38, Gencode version 31.

The Rpackage Seurat (v.4) in the R environment v.4.0.3 was used for analysis. For each sample, we dropped out cells associated with less than 200 or more than 7000 genes and having a percentage of mitochondrial genes higher than 25%. Highly variable genes were identified as the top 2000 most variable genes according to the VST transformation, corresponding to ∼8-12% of the total number of expressed genes, depending on the considered sample. The effect of the cell cycle was evaluated with a PCA and, if a relationship between cycle phases and cell segregation was observed, expression data were adjusted with a regression on cell cycle scores. Dimensional reduction by PCA was followed by UMAP algorithm, to identify similar expression profiles, and SNN unsupervised clustering algorithm, to further distinguish clusters based on top gene expression. Clustering analysis was performed considering the first 30 principal components (PCs) with resolution of 0.5. The top 50 genes for each cluster were used to infer the cluster cell type, by comparing to current single cell pancreas transcriptomes [23, 24].

To maximize cluster homogeneity among the two genotypes, we derived a reference by integrating both WFS1 and WFS1^wt/757A>T^ iBeta and performing canonical correlation analyses. Differential gene expression (DEG) analysis was carried out within each cluster setting a log2FC threshold greater than 0.25 and an adjusted p-value (FDR) lower than 0.01. Features plots using UMAP plot and violin plots reporting log-normalized UMI were used to visualize gene expression across different subpopulations and differences in gene expression between samples, respectively.

### Dynamic perifusion

Dynamic secretagogue stimulation of iPSC-derived iBeta was performed using an automated perifusion system (BioRep® Perifusion V2.0.0). Two hundred absolute clusters per condition were picked for dynamic perifusion and resuspended in the HEPES-buffered solution (125 mM NaCl, 5.9 mM KCl, 2.56 mM CaCl_2_, 1 mM MgCl_2_, 25 mM HEPES, 0.1% BSA, pH 7.4), which was supplemented as follow: 0.5 mM glucose, 11 mM glucose + 50 µM IBMX (Gibco), 11 mM glucose + 1 μM liraglutide (DBA) or 30 mM KCl. Perifusates were collected every minute; representative time points were quantified for insulin and proinsulin content by Ultrasentitive Insulin ELISA kit (Mercordia) and Human Proinsulin ELISA kit (RayBiotech).

### Ca^2+^ flux assay

Hand-picked insulin-positive clusters were plated in Optical bottom plates, 96-well (Greiner) coated with Matrigel® Basement Membrane Matrix (Corning) and they were allowed to adhere for two to three days before analysis. On the day of Ca^2+^ measuring, cells were loaded with 1 µM Fluo-4, AM, cell permeant (Invitrogen) diluted in HEPES-buffered solution for 30’ at 37°C. All image sequences were acquired at Widefield Zeiss Axio Observer.Z1. Fluo-4 was excited by a 488 laser and fluorescence signal was recorded every 2 seconds. After one minute from the start of the acquisition, cells were challenged with secretagogues and their response was recorded up to 180 seconds. Images were analyzed using the Fiji software v.1.52p and trace segments for each condition were analyzed using Matlab and R. We selected a total of 20 responder cells from two independent experiments of each condition, by drawing the region of interest (ROI) inside the cytoplasmic region of the cell. The mean of fluorescence was normalized to the corresponding mean fluorescence value of the acclimation period (F0). The change in fluorescence ΔF/F0 = (F-F0)/F0 was plotted as a function of time and cells were considered as responders if ΔF/F0 after stimulation was higher than two standard deviations from F0. Images were analyzed following background subtraction and on the resulting trace segments Matlab/R script determined number of peaks in active phase, oscillation peak amplitude, period (the length of time between two peaks), active duration A_0_ (the length of time for each Ca^2+^ oscillation measured at half of the peak height, referred to as active duration), silent duration S_0_ (expressed as the difference between period and active duration), plateau fraction (expressed as the ratio between active duration and period) and co-activity coefficient (the level of synchronization among β cells), as previously described [25, 26].

### Western blot

Pellets from at least 10^6^ cells were lysed in the M-PER™ Mammalian Protein Extraction Reagent (Thermo Fisher) and proteins were quantified using Pierce™ Rapid Gold BCA Protein Assay Kit (Thermo Fisher). Proteins were held at 95°C for 10 min, resolved on Novex™ WedgeWell™ Tris-Glycine (4-20%) and electro-transferred onto polyvinylidene difluoride membrane (PVDF) from Thermo Fisher. Transfer efficiency was evaluated using Ponceau S (Sigma). Aspecific binding sites were saturated with blocking buffer (TBS, 0.1% Tween 20 and 5% skim milk) for 1h at room temperature. Primary antibodies, which are reported in **ESM Table 1**, were incubated overnight at 4°C diluted in TBS supplemented with 0.1% Tween 20 and either 5% milk or 5% BSA. Horseradish peroxidase-conjugated secondary antibodies diluted 1:1000 in the same diluent as the corresponding primary antibody were incubated for 1h at room temperature. SuperSignal™ West Pico PLUS (Thermo Fisher) substrate was used for chemiluminescence acquisition at ChemiDoc MP (Biorad). Quantification is expressed as the mean gray area of each band and was performed using Fiji software.

### Statistical analysis

GraphPad Prism software (9.0.1 version) was employed to perform statistical analyses. Student’s unpaired or paired t-test or Mann-Whitney nonparametric test were used for the comparison of two groups and single variables. The two-way ANOVA following Dunn-Šídák’s correction for multiple comparisons was applied. In all cases, a p-value below 0.05 was considered statistically significant. Data are graphed as mean ± standard deviation (SD) unless otherwise specified. Error bar meaning, number and characteristics of replicates, and statistical analysis applied are all reported in the figure legends.

## Results

### WFS1 iPSCs efficiently differentiate *in vitro* into the pancreatic lineage, but show less mature endocrine phenotype and impaired insulin secretion

In our previous study on the molecular characterization of the c.316-1G>A mutation in the *WFS1* gene, we generated patient-derived iPSCs (WFS1) and the isogenic line in which the c.316-1G>A mutation was genetically corrected (WFS1^wt/757A>T^), restoring the wild type (WT) isoform of Wolframin ([21] and **Fig. 1a**). The mutated (WFS1) and gene edited (WFS1^wt/757A>T^) iPSC lines were differentiated into β cells (iBeta), following an *in vitro* protocol mimicking the embryonic pancreas development (**Fig. 1b** and ***Material and Methods***), to evaluate the impact of *WFS1* mutations on differentiation efficiency.

**Figure 1.**
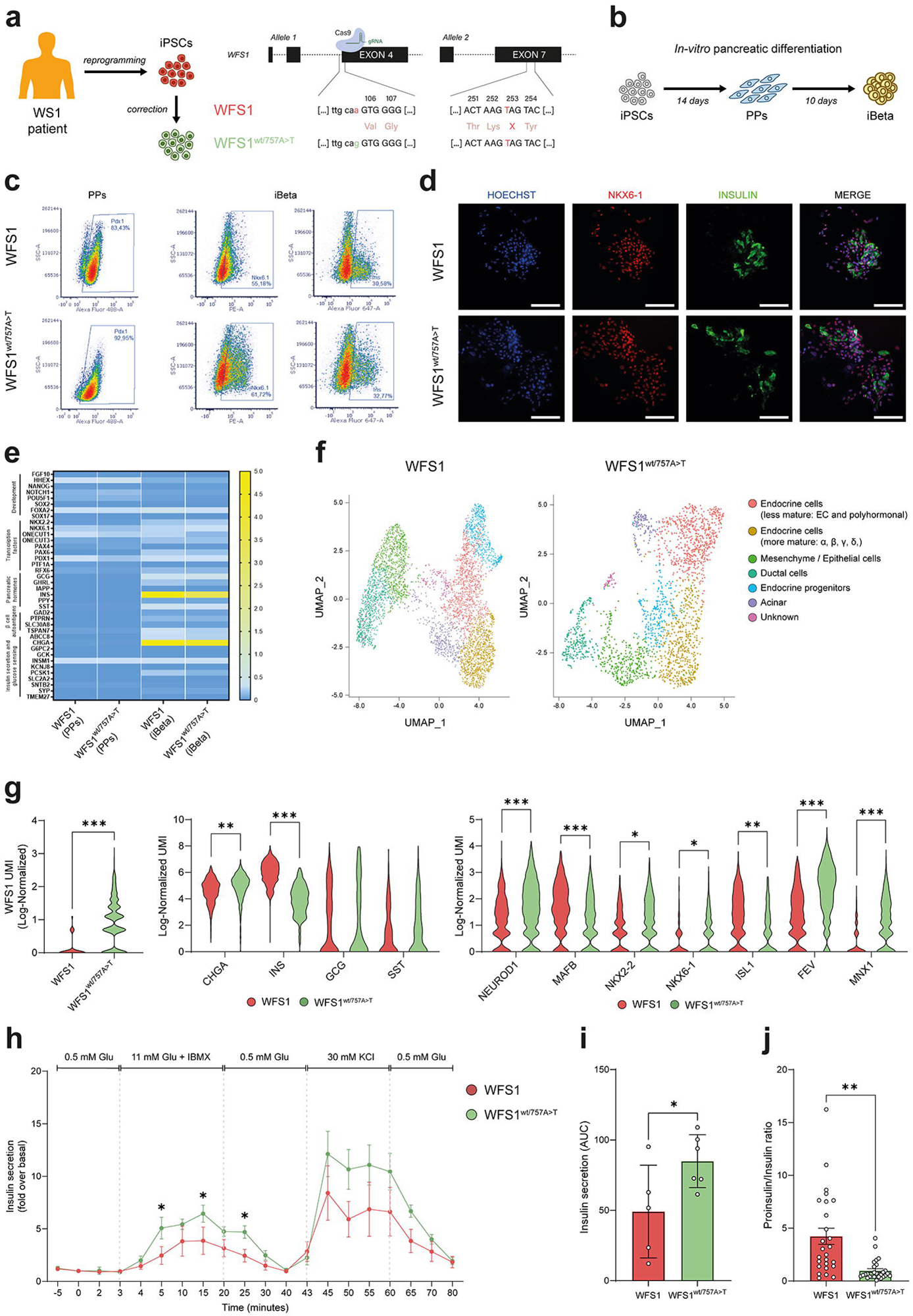
**(a)** Schematic representation of iPSCs generation from a patient with WS1 and gene targeting for CRISPR/Cas9 correction of the c.316G>A-carrying allele. **(b)** Schematic of differentiation protocol mimicking development of pancreatic progenitors (PPs) and pancreatic β cells (iBeta). **(c)** Representative flow cytometry dot plots showing the percentage of PDX1^+^, NKX6-1^+^ and Insulin^+^ cells in WFS1 and WFS1^wt/757A>T^ at the PP and iBeta stages. **(d)** Immunostaining of WFS1 and WFS1^wt/757A>T^ iBeta stained for NKX6-1 and Insulin. Nuclei were counterstained with Hoechst. Scale bar, 100 μm. **(e)** Heatmap visualization of RT-qPCR analysis of mRNA levels of endocrine markers, β cell autoantigens, insulin secretion and glucose sensing genes in WFS1 and WFS1^wt/757A>T^ at PP and iBeta stages of pancreatic development. **(f)** UMAP projection from unsupervised clustering of transcriptional data from scRNAseq of WFS1 and WFS1^wt/757A>T^ iBeta. **(g)** Violin plot detailing log-normalized gene expression of *WFS1*, *CHGA*, pancreatic hormones (*INS*, *GCG* and *SST*) and β cell maturation markers in WFS1 (red) and WFS1^wt/757A>T^ (green) endocrine clusters. *p < 0.05, **p < 0.01, ***p < 0.001 by Man-Whitney nonparametric test. **(h)** Dynamic human insulin secretion expressed as fold change over basal secretion in 0.5 mM glucose. N=5 for WFS1 and N=6 for WFS1^wt/757A>T^. *p<0.05 by unpaired two-tailed t-test. **(i)** Area under the curve (AUC) quantification of insulin secreted when clusters were perfused with 11 mM glucose + IBMX. Data are represented as mean±SD. N=5-6. *p<0.05 by unpaired one-tailed t-test. **(j)** Proinsulin/insulin ratio when clusters were perfused with 11 mM glucose. Data are represented as mean±SEM. N=15 on 3 independent experiments. **p<0.01 by unpaired one-tailed t-test.

However, as reported in the previous study [21] and confirmed here by analyzing the expression of PDX1 at the Pancreatic Progenitors (PPs) stage (**Fig. 1c**) and of NKX6-1 and Insulin at the end of differentiation (**Fig. 1c-d**), no evident defects in the pancreatic differentiation capacity were detected in WFS1 cells in comparison with the genetically correct counterpart. No significant difference between WFS1 and WFS1^wt/757A>T^ iBeta was also found by carrying out TaqMan Array microfluidic card bulk analysis of expression of the genes involved in pancreatic development, in the production of hormones and autoantigens of β cells, and in the insulin secretion and glucose sensing (**Fig. 1e**).

To overcome limitations of a bulk analysis and to highlight differences in the transcriptional profile of the endocrine compartment of differentiated cells, the expression was typified at the single cell level by performing single cell RNA sequencing (scRNAseq) on WFS1 and WFS1^wt/757A>T^ iBeta using droplet-based Chromium 10X platform and Illumina NovaSeq. After quality control, the effect of cell cycle was removed from the expression data by using a linear regression model and we maintained for further analysis a total of 4,594 and 2,215 iBeta cells differentiated from WFS1 and WFS1^wt/757A>T^ iPSCs, respectively (**ESM Fig. 1a-f**). The unsupervised clustering was performed to identify a total of 7 subpopulations, based on the top upregulated genes (**ESM Fig. 1g-h**). Cluster annotation was obtained by matching upregulated genes of each cluster to the published transcriptome data of human pancreatic cell types (**Fig. 1f**) [23, 24].

According to our previous investigations, we found lower expression of Wolframin in WFS1 iBeta, consisting of the residual stable internally-truncated isoforms (**Fig. 1g**). Interestingly, WFS1 iBeta displayed significantly higher expression of *INS* (5.92±0.88 vs 3.90±1.13; p < 0.001) in comparison to WFS1^wt/757A>T^, but a slight reduction of *CHGA* (4.48±0.90 vs 4.89±1.02; p < 0.01). Although the average expression levels of *GCG* and *SST* were comparable (**Fig. 1g**), the detailed analysis of the subpopulations within the endocrine compartments highlighted strong differences between the two samples. Compared to WFS1^wt/757A>T^, WFS1 iBeta had a greater proportion of β cells (85.9% vs 75.7%), but a significantly lower one of mature α (1.4% vs 12.7%), δ (0.2% vs 4%) and PP (0% vs 0.9%) cells (**ESM Fig. 2a-b**). By comparing the respective endocrine clusters for the expression levels of transcription factors involved in endocrine commitment and β cell maturation, we found that WFS1 cells displayed significantly higher levels of the early endocrine markers *MAFB* and *ISL1*, but a reduced expression of *NEUROD1*, *NKX2-2*, *NKX6-1* and *FEV* in comparison to WFS1^wt/757A>T^. Furthermore, the WFS1 endocrine cells showed a FEV-to-ISL1 ratio ≈ 1, whereas in WFS1^wt/757A>T^ it was > 2 (**Fig. 1g**). Consistent with the evidence that the onset of FEV preceding ISL1 initiation is indicative of α/β cell fate and, vice versa, the expression of ISL1 over FEV appears to predict an immature, polyhormonal profile [27], we found reduced expression in WFS1 iBeta of *MNX1* (0.37±0.57 vs 1.21±0.87; p<0.001), a transcription factor pivotal for β cell-like maturation (**Fig. 1g**), and more INS^+^/GCG^+^/SST^+^ or INS^+^/GCG^+^/PPY^+^ polyhormonal cells in WFS1 (12.2%) compared to WFS1^wt/757A>T^ (6.4%) iBeta (**ESM Fig. 2a-b**). Further evaluation of the late β cell markers we found differentially expressed between two lines, such as *UCN3*, *SYT4*, *CFAP126* and *MAFA*, supported the hypothesis that WFS1 iBeta displayed a less mature phenotype than its gene edited counterpart (**ESM Fig. 2c-d**). Finally, after considering the maturation differences of INS^+^ cells between the two iBeta lines, we found that WFS1 cells exhibited a reduced glucose responsiveness in dynamic perifusion and a significantly lower (49.09±32.92 vs 84.93±18-81; p<0.05) insulin secretion area under curve (AUC) than their gene edited counterpart (**Fig. 1h-i**). The proinsulin/insulin ratio, calculated during the 11 mM glucose stimulus step of dynamic perifusion, was also altered in WFS1 cells compared to the gene edited cell line (4.12±0.95 vs 0.69±0.14; p<0.01), suggesting a disruption in the normal processing of insulin in mutated cells, reflecting underlying β cell stress or damage (**Fig. 1j**).

**Figure 2.**
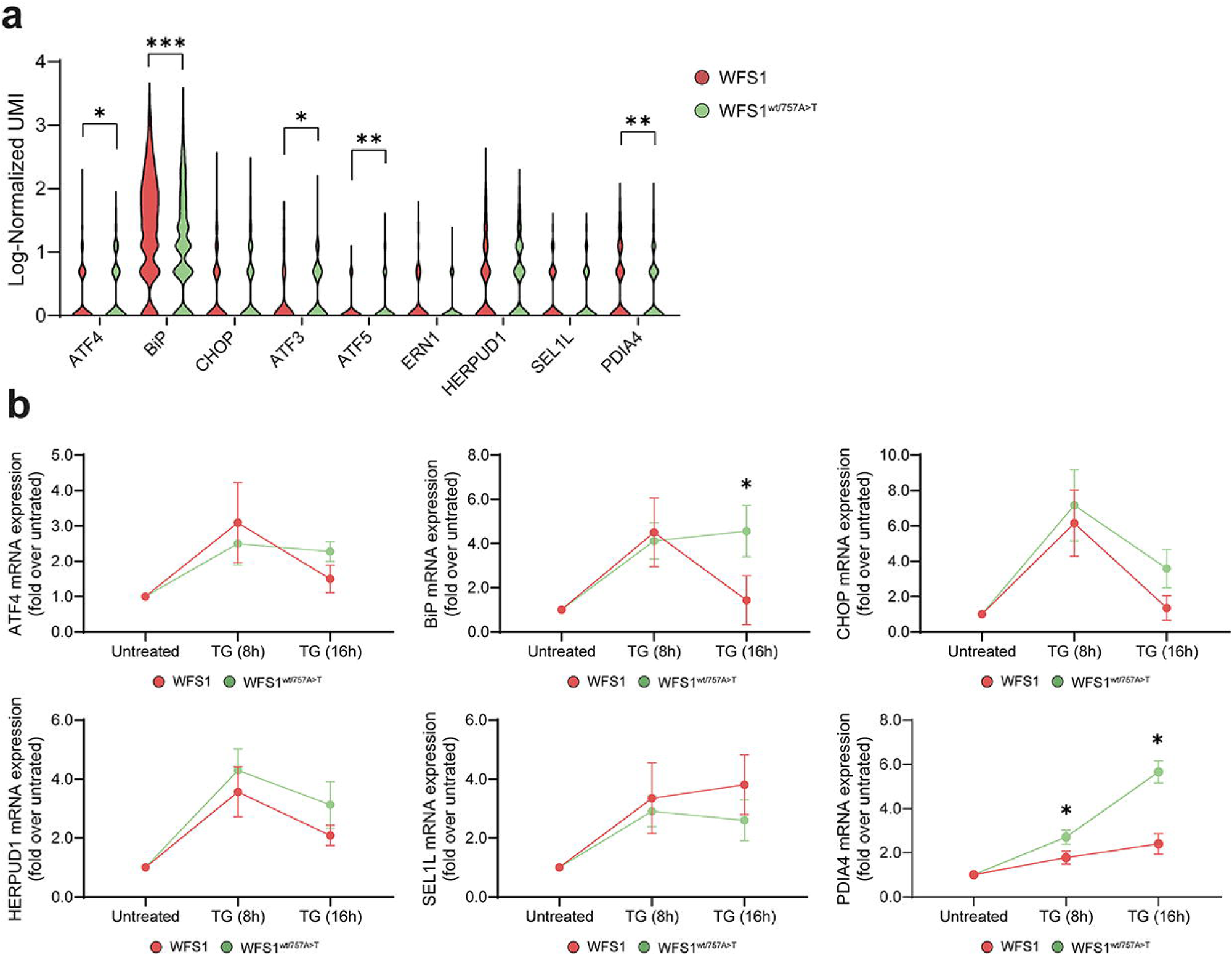
**(a)** Violin plot reporting the log-normalized expression of ER-stress related genes in the endocrine clusters of WFS1 (red) and WFS1^wt/757A>T^ (green) iBeta. *p < 0.05, **p < 0.01, ***p < 0.001 by Man-Whitney nonparametric test. **(b)** Fold increase of ER-stress markers *ATF4*, *BiP*, *CHOP*, *HERPUD1*, *SELS1L* and *PDIA4* in WFS1 and WFS1^wt/757A>T^ iBeta after 50 nM TG treatment at the indicated times. Data are plotted as mean±SD. N=7-9. *p<0.05 by unpaired two-tailed t-test.

### WFS1 iBeta have negligible impairment in ER stress-related markers under basal conditions, but an inadequate UPR upon ER stress induction

As pathogenic variants in Wolframin cause ER stress in β cells, we investigated the expression of ER stress-related markers in the endocrine compartment of WFS1 iBeta in comparison to the syngeneic gene edited cells. However, as previously observed by performing both RT-qPCR and immunoblot analysis on bulk samples [21], we did not find a general increase in stress markers in WFS1 endocrine cells. With the exception of the ER stress chaperones BiP (1.40±0.81 in WFS1 vs 1.11±0.80 in WFS1^wt/757A>T^) and PDIA4 (0.43±0.53 in WFS1 vs 0.35±0.48 in WFS1^wt/757A>T^), we did not find differences in several ER stress markers, including CHOP, ERN1 and SEL1L, whereas we noted a significant reduction of ATF3, ATF4 and ATF5 in WFS1 compared to WFS1^wt/757A>T^ (**Fig. 2a**).

To further investigate the effects of cellular stress, we treated WFS1 and WFS1^wt/757A>T^ with 50 nM TG for 8h and 16h, to induce UPR activation. As we previously highlighted [21], the response during the first hours of treatment appears overall comparable between the two cell lines (**Fig. 2b**), confirming the functioning of the residual Wolframin isoforms, whose expression was significantly induced following TG treatment (**ESM Fig. 3**). However, prolonged ER stress (∼16h) highlighted UPR impairments in WFS1 iBeta and a significant decrease of the chaperones BiP and PDIA4 compared to WFS1^wt/757A>T^ (**Fig. 2b**), suggesting that WFS1 cells were unable to sustain long-term response to cellular stress and to escape from apoptosis [21].

**Figure 3.**
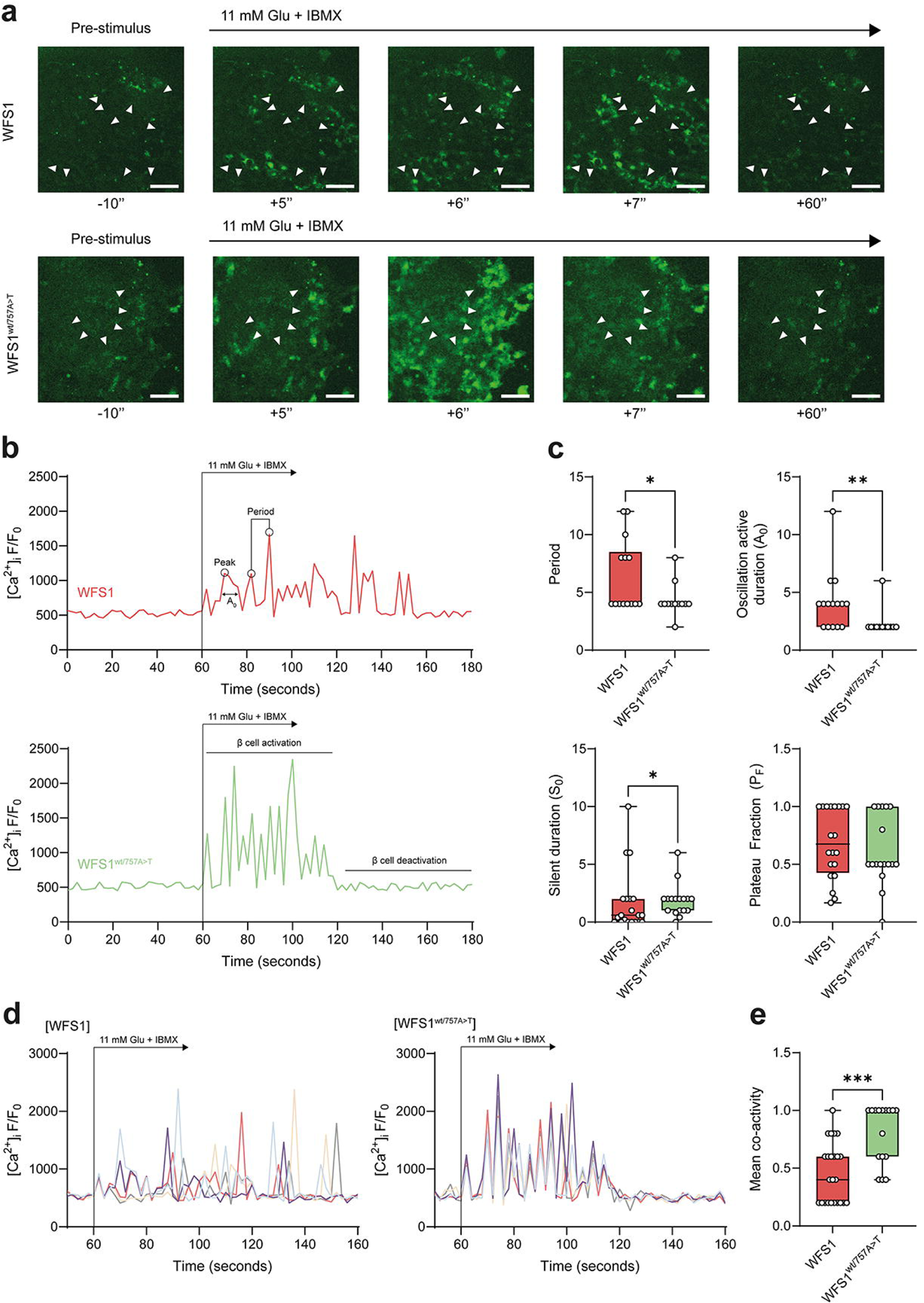
**(a)** Imaging of Ca^2+^ flux using Fluo-4 fluorescent dye in WFS1 and WFS1^wt/757A>T^ β cells before (-10 seconds) and after (+5, 6, 7 and 60 seconds) stimulation with 11 mM glucose + IBMX. White triangles indicate responding cells. Scale bar, 50 μm. **(b)** Mean fluorescence signal (F/F_0_) over time in WFS1 (red) and WFS1^wt/757A>T^ (green) iBeta indicating [Ca^2+^]_i_ elevation in response to stimulation with 11 mM glucose + IBMX. Peak (the maximum value of each oscillation), period (the length of time between two peaks) and active duration (A_0_ - the length of time for Ca^2+^ oscillation measured at half of the peak height) are indicated within the WFS1 (red) graph. The timing of β cell activation and de-activation are highlighted in the WFS1^wt/757A>T^ (green) graph. N=20 cells from 2 independent experiments. **(c)** Mean period upon stimulation, active and silent duration of oscillation, and plateau fraction measured in WFS1 and WFS1^wt/757A>T^ β cells. N=20. *p<0.05, **p<0.01 by Man-Whitney nonparametric test. **(d)** Five typical [Ca^2+^]_i_ traces recorded during stimulation with 11 mM glucose + IBMX of a single cluster of WFS1 and WFS1^wt/757A>T^ β cells. **(e)** Mean coactivity coefficient pooled from different clusters of WFS1 and WFS1^wt/757A>T^. N=20 cells from 2 independent experiments. ***p<0.001 by Mann-Whitney nonparametric test.

### WFS1 iBeta displays altered Ca^2+^ dynamics and dysregulation of insulin secretion machinery

To deeply understand the mechanisms driving the impairment of the insulin secretion in WFS1 iBeta, we measured the Ca^2+^ dynamics in response to secretagogue stimulation. As reported in **Fig. 3a-c**, multiple properties of Ca^2+^ fluxes were different between WFS1 and WFS1^wt/757A>T^ iBeta. Indeed, while Ca^2+^ concentrations oscillated synchronously in WFS1^wt/757A>T^ iBeta, as demonstrated by the levels of Fluo-4 during live-cell imaging in response to glucose stimulation, the Ca^2+^ waves did not propagate simultaneously through the cell population in WFS1 iBeta, which appeared to be also affected by delayed β deactivation (**Fig. 3a-b**). To better elucidate differences between the two iBeta, we measured key parameters of the Ca^2+^ waveform. The transition from low to high glucose concentration resulted in oscillations that changed in their average peak and showed increased period (the length of time between two peaks) and time in the active phase A_0_ (the length of time for each Ca^2+^ oscillation measured at half of the peak height, referred to as active duration) in WFS1 iBeta compared to the WFS1^wt/757A>T^ counterpart. Conversely, the silent duration S_0_ (the electrically silent ‘triggering’ phase, which culminates in membrane depolarization) appeared to be reduced in WFS1 iBeta, while the plateau fraction P_F_ (which measures the proportion of time spent in active phase over a single period) did not show significant differences between the two genotypes (**Fig. 3b-c**). Typical single-cell Ca^2+^ traces (**Fig. 3d)** and the mean of co-activity, calculated as the number of cells with similar peak profiles (**Fig. 3e**), support the evidence that Ca^2+^ oscillations were less synchronous in WFS1 compared to WFS1^wt/757A>T^ iBeta.

As the exocytosis of insulin-containing granules in pancreatic β cells is essential to their function, we investigated the pathways involved in the priming and fusion of secretory granules with plasma membrane preceding their Ca^2+^-dependent exocytosis and consequent insulin release [28]. During the early events, the plasma membrane associated proteins syntaxin 1A (Stx1A) and synaptosomal-protein of 25 kD (SNAP25) interact with the vesicle-associated membrane protein (Vamp2), to form the Soluble NSF-Attachment Protein (SNAP) Receptors (SNARE) complex that promotes fusion by pulling the vesicle membrane in close contact with the plasma membrane upon Ca^2+^ influx [29]. In the endocrine compartment of iPSC pancreatic derivatives, we examined the expression of *RGS4*, encoding for the regulator of G protein signaling 4 (RGP4), a negative regulator of insulin secretion in pancreatic β cells [30], and of *SNAP25* and *VAMP2* genes, as key essential component of the insulin exocytosis machinery.

We also evaluated the *CACNA1C* and *CACNA1D* genes, respectively coding for calcium voltage-gated channel subunit alpha1 C (CACNA1C, also known as Ca_v_1.2) and alpha1 D (CACNA1D, also known as Ca_v_1.3). The scRNAseq revealed an *RGS4* upregulation (1.50±1.02 in WFS1 vs 1.32±1.06 in WFS1^wt/757A>T^; p<0.01) and concomitant *CACNA1C* (0.08±0.23 in WFS1 vs 0.15±0.30 in WFS1^wt/757A>T^; p<0.001) and *CACNA1D* (0.10±0.28 in WFS1 vs 0.27±0.44 in WFS1^wt/757A>T^; p<0.001) decrease (**Fig. 4a**), indicating a strong inhibition of calcium release in WFS1 iBeta, that is at least in part due to an impairment in the Ca^2+^ channels compared to the WFS1^wt/757A>T^ counterpart. Moreover, WFS1 iBeta showed significantly reduced transcriptional levels of *SNAP25* (0.19±0.35 in WFS1 vs 0.45±0.57 in WFS1^wt/757A>T^; p<0.001) (**Fig. 4a**), suggesting a defect of the β cell secretory machinery.

**Figure 4.**
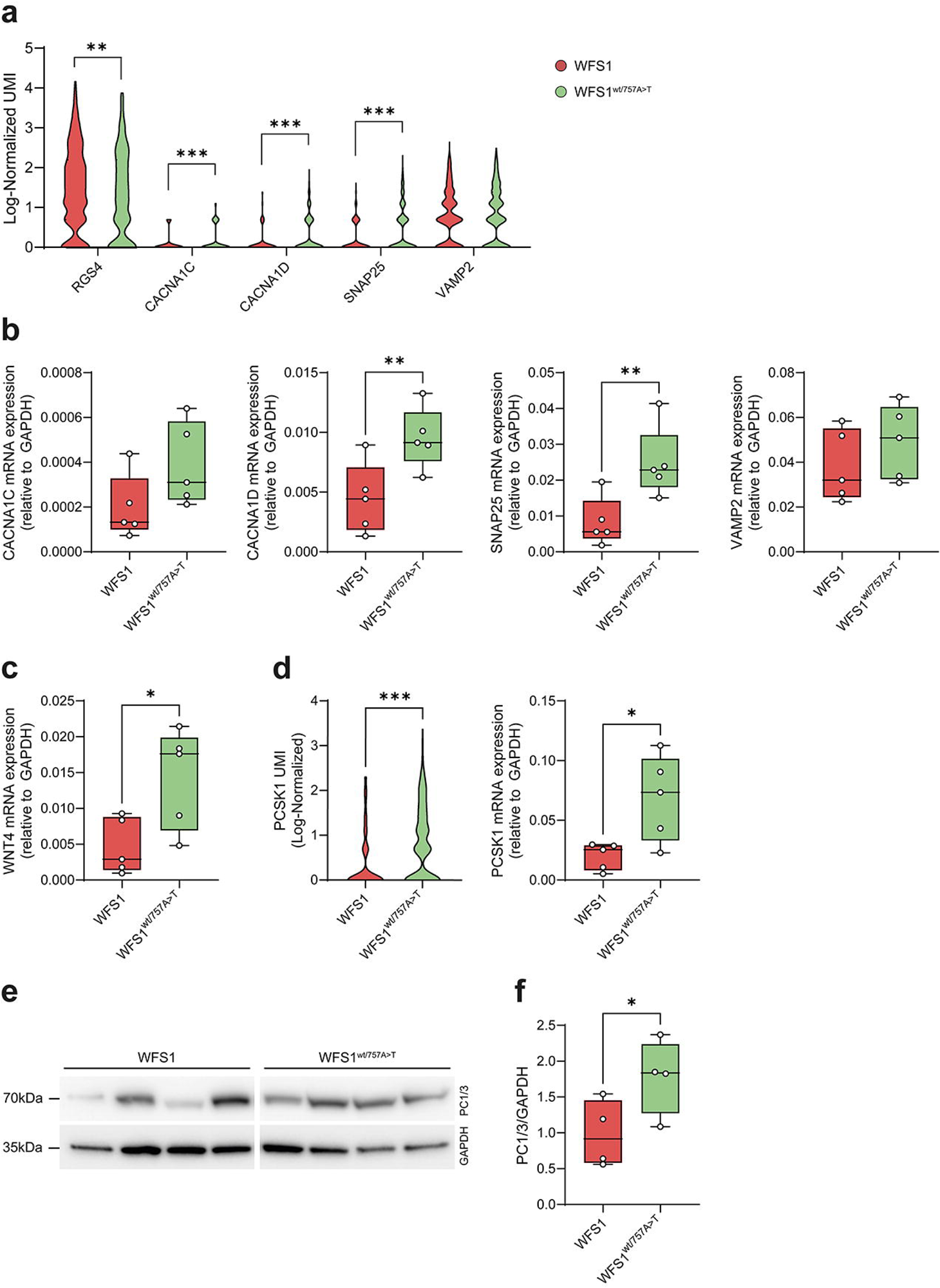
**(a)** Violin plot reporting the log-normalized expression of Ca^2+^-related (*RGS4*, *CACNA1C* and *CACNA1D*) and SNARE complex-related (*SNAP25* and *VAMP2*) genes in the endocrine clusters of WFS1 (red) and WFS1^wt/757A>T^ (green) iBeta. *p < 0.05, ***p < 0.001 by Mann-Whitney nonparametric test. The expression levels measured by RT-qPCR of **(b)** *CACNA1C-D*, *SNAP25* and *VAMP2*, and **(c)** *WNT4* genes in WFS1 (red) and WFS1^wt/757A>T^ (green) iBeta. N=5. **p < 0.001 by unpaired two-tailed t-test. **(d)** Log-normalized expression of *PCSK1* gene in the endocrine clusters from scRNAseq experiment and analysis of its expression levels by RT-qPCR in WFS1 (red) and WFS1^wt/757A>T^ (green) iBeta. ***p < 0.001 by Mann-Whitney nonparametric test (for scRNAseq); N=5. *p < 0.05 by unpaired two-tailed t-test (for RT-qPCR). **(e)** Immunoblot analysis of PC1/3 expression in WFS1 and WFS1^wt/757A>T^ iBeta. **(f)** Relative quantification of PC1/3 to GAPDH (housekeeping) expression in WFS1 and WFS1^wt/757A>T^ iBeta. N=4. *p<0.05 by unpaired one-tailed t-test.

The quantification of mRNA transcripts by RT-qPCR confirmed a statistically significant decrease in *CACNA1D* mRNA, but not *CACNA1C* (**Fig. 4b**), suggesting that Ca_v_1.3 is relevant for insulin secretion in WS1. Moreover, in WFS1 iBeta we found >2-fold lower levels of *WNT4* mRNA (0.004±0.0038 vs 0.014±0.007; p<0.05), an important factor that controls Ca^2+^ signaling in response to a glucose challenge [31], suggesting that the Wnt pathway is affected in WS1 and may contribute to further impairments in glucose-dependent insulin secretion (**Fig. 4c**). We further investigated whether insulin processing was affected in WFS1 iBeta, and we observed a reduction of *PCSK1* gene expression (**Fig. 4d**) and its protein product PC1/3 (**Fig. 4e-f**) in these cells compared to WFS1^wt/757A>T^. All together, these results highlight the dysregulation of several β cell-specific pathways that ultimately lead to impaired β cell function in WS1.

### WFS1 iBeta shows enhanced autophagic flux under basal conditions

To give a deeper insight into the mechanisms underlying the reduced insulin secretion in WFS1 iBeta, we further investigated cellular pathways potentially involved in this process. It has been reported that the loss of Wolframin and the consequent alteration of the Ca^2+^ dynamics activate autophagy and mitophagy to supply mitochondrial metabolism and increase energy production [11, 12]. Based on this evidence, we attempted to investigate autophagy in WFS1 iBeta from the patient under investigation. We first analyzed the transcription levels of the key regulators *ATG10*, *ATG12* and *BECN1* that participate in autophagosome formation, but we did not find statistically significant differences between WFS1 iBeta and their corrected counterpart, except for significantly reduced *ATG10* transcripts in WFS1 iBeta (0.0056±0.0010 vs 0.0098±0.0017) (**Fig. 5a**). Surprisingly, the autophagy substrate *SQSTM1* mRNA was strongly upregulated in WFS1 (0.24±0.10) compared to WFS1^wt/757A>T^ (0.033±0.004) iBeta (**Fig. 5a**). To detect the autophagic events, we further analyzed the accumulation of the lipidated LC3 (LC3-II) by immunoblotting and we did not find any difference in the LC3-II levels between WFS1 iBeta and WFS1^wt/757A>T^ iBeta in basal conditions. We also monitored the abundance of SQSTM1 by immunoblot and observed reduced expression levels in WFS1 compared to WFS1^wt/757A>T^ iBeta (**Fig. 5b-c**). Since the low expression levels of SQSTM1 can either result either from a reduction of autophagosome formation or from an increase in fusion with lysosomes and subsequent SQSTM1 degradation, we explored the autophagic flux by treating iBeta with bafilomycin, that prevents the acidification of lysosomes. LC3-II accumulated up to 2-fold in presence of bafilomycin in both WFS1 (p<0.01) and WFS1^wt/757A>T^ (p<0.05) iBeta, but the SQSTM1 levels increased >2.5-fold only in WFS1 iBeta treated with bafilomycin (p<0.05), suggesting that Wolframin loss enhances the autophagic flux (**Fig. 5d-e**). These results are indeed concordant with the increase of *SQSTM1* transcripts in WFS1 iBeta, which constitutes an attempt at restoring SQSTM1 protein levels.

**Figure 5.**
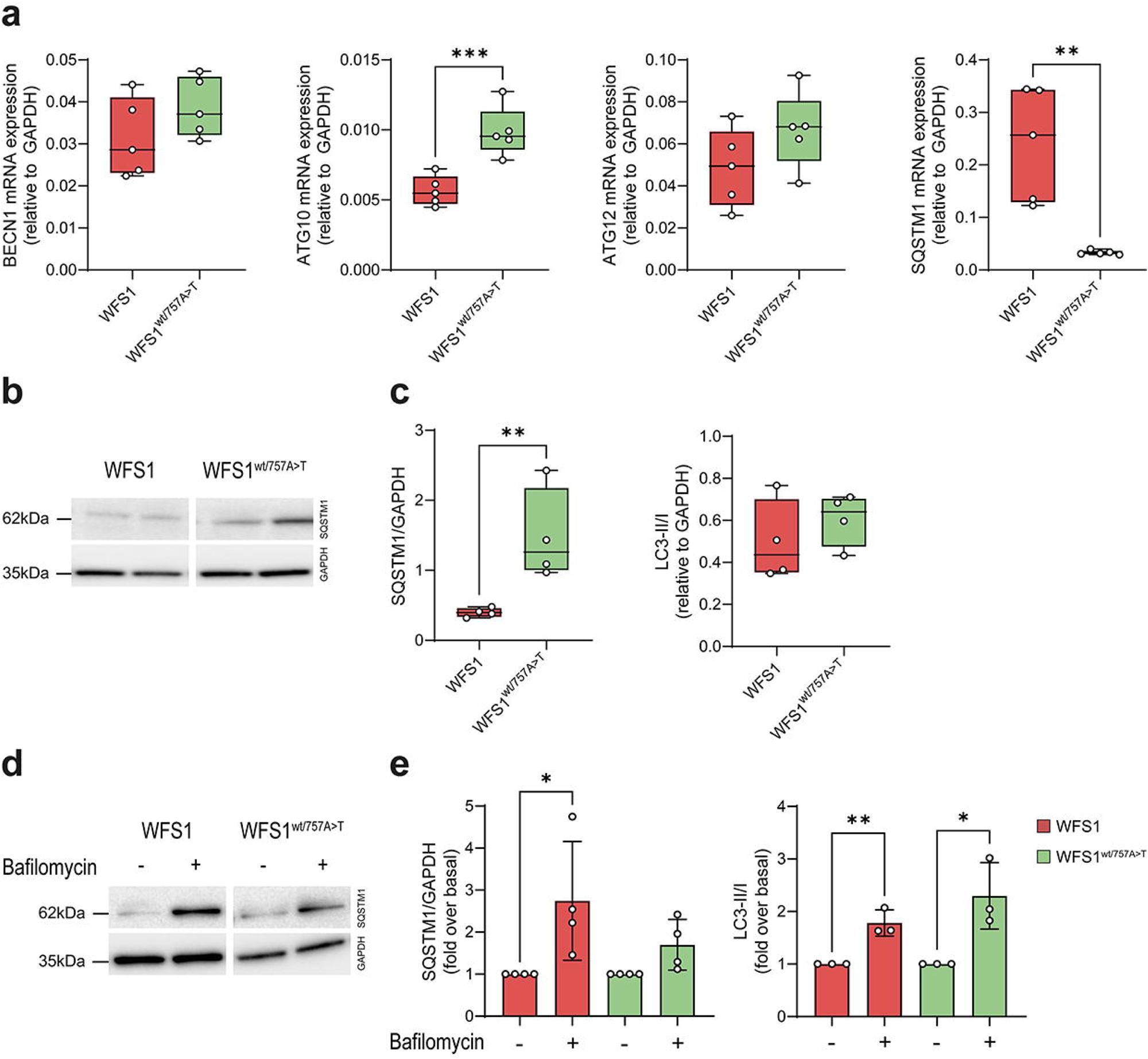
**(a)** Expression levels measured by RT-qPCR of *BECN1*, *ATG10*, *ATG12* and *SQSTM1* genes in WFS1 (red) and WFS1^wt/757A>T^ (green) iBeta. *GAPDH* was used as a housekeeping gene. N=5. **p<0.01, ***p<0.001 by unpaired two-tailed t-test. **(b)** Representative immunoblots for analysis of SQSTM1 in WFS1 and WFS1^wt/757A>T^ iBeta. **(c)** Relative quantification of the SQSTM1 protein and the LC3-II/I ratio using GAPDH as normalizer in WFS1 (red) and WFS1^wt/757A>T^ (green) iBeta. N=4. **p<0.01 by unpaired one-tailed t-test. **(d)** Immunoblot analysis of SQSTM1 expression in WFS1 and WFS1^wt/757A>T^ iBeta treated with 100 nM bafilomycin A_1_ for 24h. **(e)** Relative quantification of SQSTM1 and LC3-I/LC3-II ratio in bafilomycin A_1_-treated WFS1 and WFS1^wt/757A>T^ iBeta. GAPDH was used as housekeeping. Data are reported as mean±SD expressed as fold increase over untreated cells. N=3 independent experiments. *p<0.05, **p<0.01 by paired two-tailed t-test.

### The GLP-1R agonist liraglutide restores the Ca^2+^ oscillation and secretory impairments, increasing glucose responsiveness and insulin release

Finally, we sought to investigate whether the mechanisms we found dysregulated in patient WFS1 iBeta might be positively impacted by liraglutide treatment and, therefore, whether their targeting might account for the efficacy of liraglutide in this patient [18]. We first inspected the effect of acute liraglutide treatment on glucose responsiveness and insulin secretion in WFS1 iBeta. We observed 2-fold increase of secreted insulin in WFS1 iBeta following stimulation with 11 mM glucose and 1 μM liraglutide in comparison with cells stimulated with 11 mM glucose and IBMX (**Fig. 6a**). We quantified the magnitude of glucose responsiveness by measuring the insulin secretion AUC, which increased significantly from 49.08±32.9 in untreated WFS1 iBeta to 84.68±14.11 (p<0.05) after liraglutide treatment (**Fig. 6b**). Given the improvement observed on insulin secretion capacity, we evaluated if liraglutide-related effects correlated with overall amelioration of Ca^2+^-related parameters. As shown in **Fig. 6c-d**, liraglutide was able to modify the Ca^2+^ oscillation of WFS1 iBeta upon stimulation, resulting in increased average peak intensity (1118 for liraglutide-treated iBeta vs 823 for untreated iBeta), anticipated de-activation phase (start at 58’ upon stimulus for liraglutide-treated iBeta and at 94’ upon stimulus for untreated iBeta) and overall synchronization degree at comparable levels to those observed in the WFS1^wt/757A>T^ iBeta (**Fig. 6c-d**). While the period did not significantly change between the two experimental groups, the active phase of oscillations was lower in the liraglutide-treated iBeta (2.66±1.17 vs 4.13±2.55; p<0.05) and consistently the silent period of oscillations significantly increased (2.28±1.13 vs 1.67±2.66; p<0.01) (**Fig. 6e**). Accordingly, the mean of co-activity rose to 67% after liraglutide exposure in comparison to that observed in untreated cells (**Fig. 6e**). Interestingly, the acute administration of liraglutide ameliorated average peak and Ca^2+^ oscillations in both WFS1 and WFS1^wt/757A>T^ iBeta treated with 50 nM TG for 8h (**ESM Fig. 4a-b**), supporting the hypothesis that GLP-1R agonists can actively counteract cellular stress-dependent impairments. In light of this, we found that liraglutide increased the transcription of both *SNAP25* (0.048±0.019 vs 0.023±0.011; p<0.05) and *CACNA1D* (0.0083±0.0014 vs 0.0052±0.0014; p<0.05), suggesting that the exocytosis machinery as well as the calcium voltage-gated channel could be its pharmacological targets. Conversely, the drug did not seem to have any significant impact on *VAMP2* and on the autophagy substrate *SQSTM1* expression (**Fig. 6f**).

**Figure 6.**
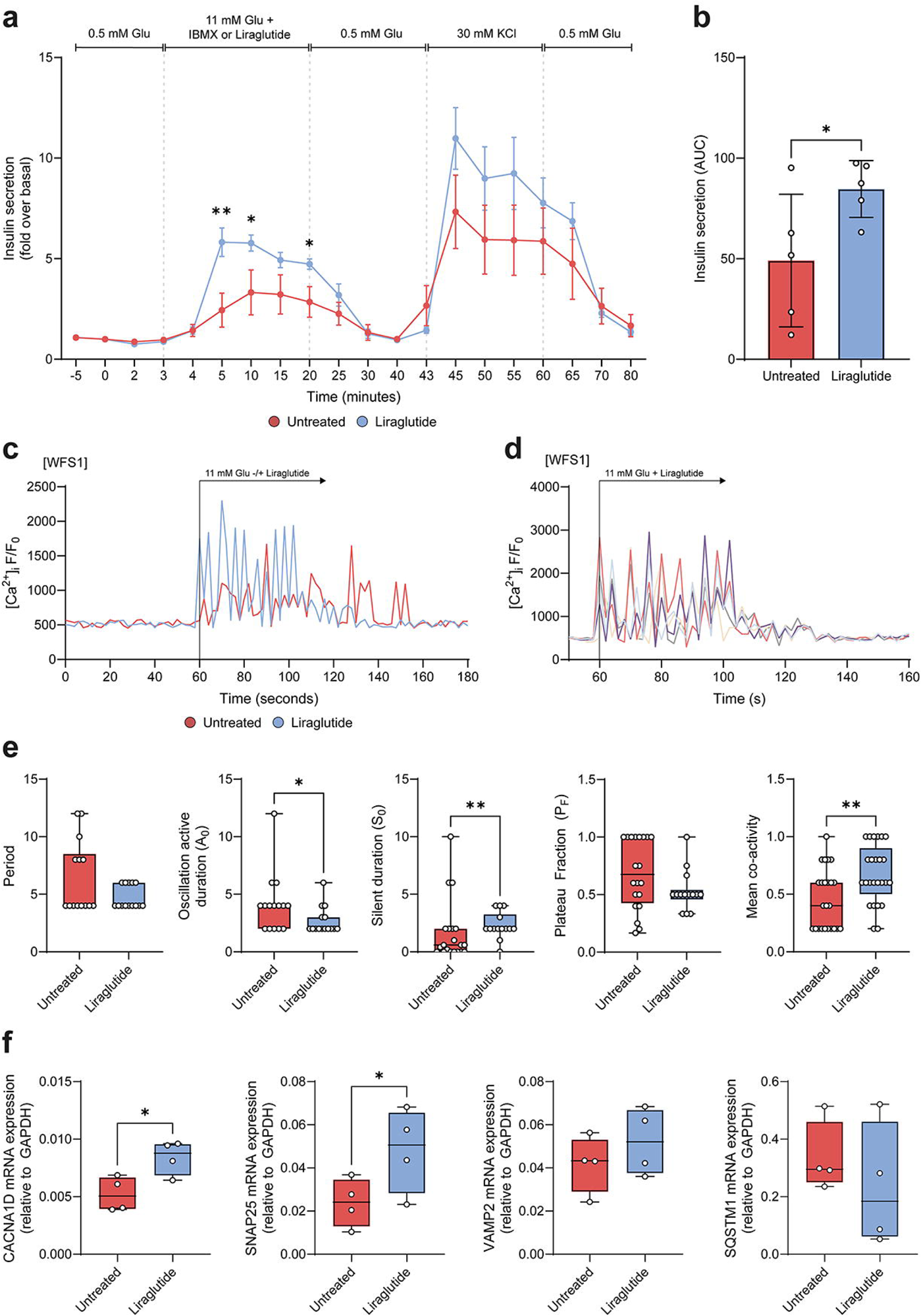
**(a)** Dynamic human insulin secretion expressed as fold change over basal secretion of WFS1 iBeta following stimulation with 11 mM glucose plus either IBMX (red) or liraglutide (blue). N=5. *p<0.05, **p<0.01 by unpaired two-tailed t-test. (**b)** Area under the curve (AUC) quantification of insulin secreted when clusters were perfused with 11 mM glucose plus either IBMX (red) or liraglutide (blue). Data are represented as mean±SD. N=5. *p<0.05 by unpaired one-tailed t-test. **(c)** Mean fluorescence signals (F/F_0_) over time indicating [Ca^2+^]_i_ oscillations in WFS1 β cells following 11 mM glucose stimulation without (red) or with (blue) liraglutide. N=20 from 2 independent experiments. **(d)** Five typical [Ca^2+^]_i_ traces recorded during stimulation with 11 mM glucose + liraglutide of WFS1 β cells. **(e)** Mean period, active and silent duration of oscillations, plateau fraction and mean co-activity coefficient in WFS1 β cells following 11 mM glucose stimulation without (red) or with (blue) liraglutide. N=20. *p<0.05, **p<0.01 by Mann-Whitney nonparametric test. **(f)** RT-qPCR analysis of the expression levels of *CACNA1D*, the SNARE complex-related genes *SNAP25* and *VAMP2*, and *SQSTM1* in WFS1 iBeta before and after treatment with liraglutide. N=4. *p<0.05 by unpaired one-tailed t-test.

### Chronic liraglutide exposure protects WFS1 iBeta from TG and pro-inflammatory cytokines-induced apoptosis

In our previous work we determined that WFS1 iBeta are more susceptible to cellular stress and to pro-inflammatory cytokines-induced cell death [21]. To uncover a possible protective role of liraglutide treatment, we evaluated the effect of prolonged exposure on cell survival under stress or inflammatory conditions (**Fig. 7a**). Strikingly, by adding liraglutide to culture medium during either TG or cytokine treatment, we observed that WFS1 iBeta were protected from undergoing both early and late apoptosis, as the percentage of both Annexin V^+^ and PI^+^ cells were comparable to the untreated ones and >2-fold lower than WFS1 iBeta treated with TG for 16h or pro-inflammatory cytokines for 48h (**Fig. 7b-d**).

**Figure 7.**
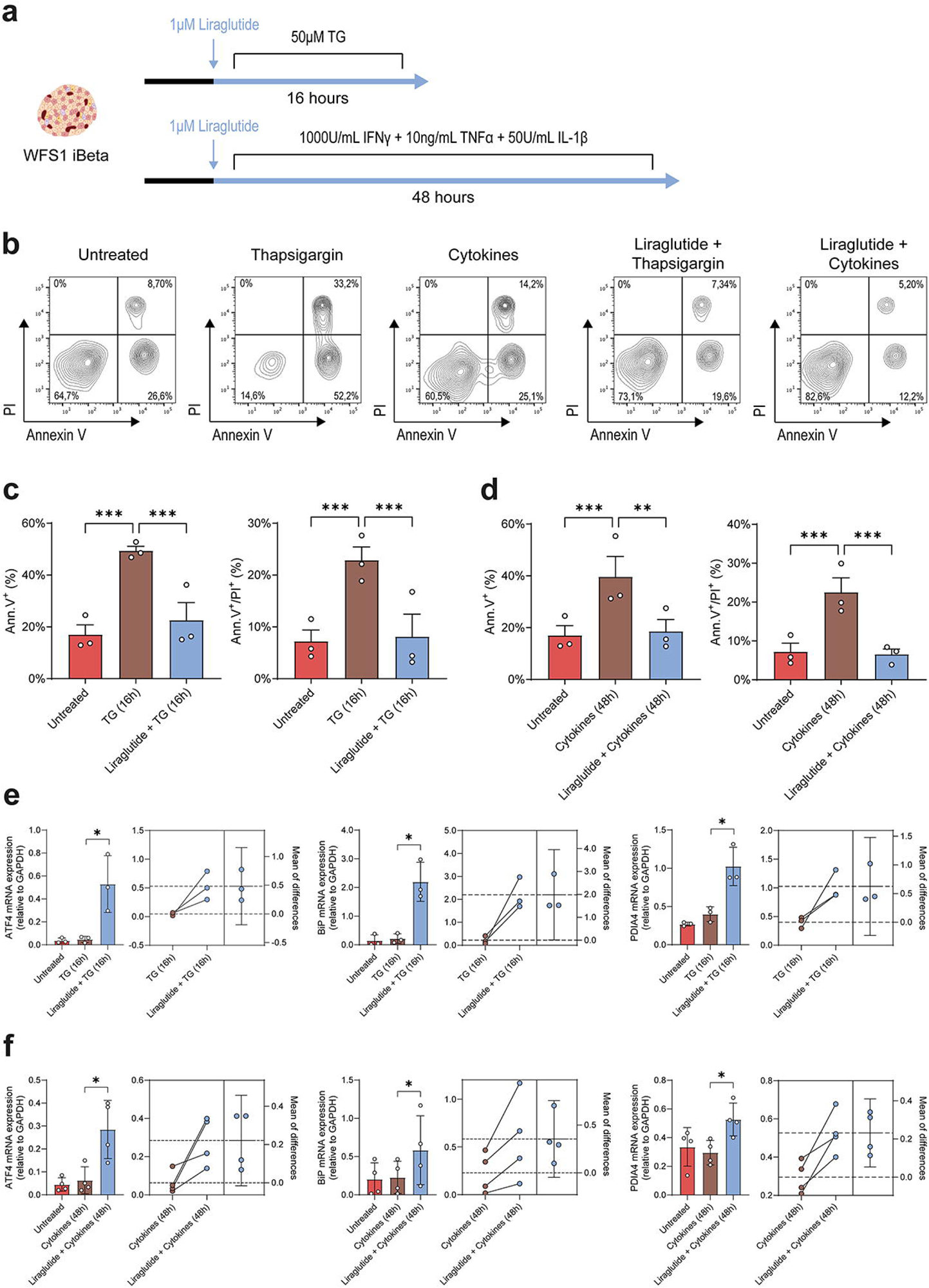
**(a)** Schematic summary of chronic treatment with 1 μM liraglutide in WFS1 iBeta exposed either to TG for 16h, or inflammatory cytokines (IFNγ, TNFα and IL-1β) for 48h. **(b)** Representative FACS plots for analysis of Annexin V (Ann.V) and PI-stained WFS1 iBeta after treatment with either TG or cytokines, in presence or absence of liraglutide. Early and late apoptosis were measured as percentage (%) of Ann.V and PI-positive cells, respectively, in WFS1 iBeta after exposure to **(c)** TG or **(d)** cytokines ± chronic treatment with liraglutide. Data are expressed as mean±SD. N=3. **p<0.01, ***p<0.001 by two-way ANOVA with Dunn-Šídák correction. **(e)** Relative gene expression of *ATF4*, *BiP* and *PDIA4* in WFS1 iBeta after 16h post TG exposure ± liraglutide, and estimation plots including the mean of difference of *ATF4*, *BiP* and *PDIA4* mRNAs following chronic treatment with liraglutide. Data are plotted as mean±SD. N=3. *p<0.05 by one-tailed unpaired t-test. **(f)** Relative gene expression of *ATF4*, *BiP* and *PDIA4* in WFS1 iBeta after 48h post pro-inflammatory cytokine exposure ± liraglutide, and estimation plots including the mean of difference of *ATF4*, *BiP* and *PDIA4* mRNAs following chronic treatment with liraglutide. Data are plotted as mean±SD. N=4. *p<0.05 by one-tailed unpaired t-test.

The capacity of liraglutide to mitigate apoptosis correlated with enhanced UPR in treated WFS1 iBeta. By evaluating the expression changes of ER stress-related markers, we found a significant increase of ATF4 (mean diff. 0.48±0.27; p<0.05), BiP (mean diff. 1.92±0.79; p<0.05) and PDIA4 (mean diff. 0.62±0.34; p<0.05) in WFS1 cells exposed to liraglutide and TG (**Fig. 7e**), and of ATF4 (mean diff. 0.22±0.14; p<0.05), BiP (mean diff. 0.35±0.25; p<0.05) and PDIA4 (mean diff. 0.23±0.11; p<0.05) in WFS1 cells exposed to liraglutide and pro-inflammatory cytokines (**Fig. 7f**).

## Discussion

Wolframin is a transmembrane ER multifunctional protein expressed in pancreatic β cells, brain, heart, lung. ER functions as a calcium storage and a quality control system driving the degradation of proteins with altered conformation and interacts with mitochondria to orchestrate Ca^2+^ and lipids transfer and regulate apoptosis and autophagy [32–34]. Wolframin is involved in the cross-talk of several signaling pathways as many studies unveiled its role in ER stress response, protein misfolding control, regulation of intracellular Ca^2+^ homeostasis [6]; moreover, it participates to the complex ER/mitochondria interplay as it takes part in mitochondria-associated ER membranes (MAMs) [34]. The critical impact of this protein on multiple cellular processes explains the clinical complexity of WS1, encompassing diabetes mellitus, optic atrophy, deafness and neurological disorders. The heterogeneity of clinical manifestations mirrors the spectrum of mutations of the *WFS1* gene, including missense, non-sense, and frameshift insertion or deletion mutations, that have dramatically different effects at both RNA and protein levels [35, 36].

We investigated a WS1 patient carrying the heterozygous c.316-1G>A mutation in the *WFS1* gene, affecting the acceptor splice site upstream exon 4, that originates two PTC-containing alternative splicing transcripts (c.316del and c.316-460del), and two ORF-conserving mRNAs (c.271-513del and c.316-456del). While the first two mRNAs are regulated by NMD and undergo degradation, the others are preserved and translated onto N-terminally truncated polypeptides, that retained the functional C-terminal domain [21]. This peculiar transcriptional landscape conferred an unconventional cellular response to the induction of ER stress in iPSC-derived b cells, that drive us to in depth characterize the physiological impact of this mutation, from the differentiation of patient-derived iPSCs into the pancreatic lineage to their capacity to respond to glucose stimulation.

Indeed, β cell functional maturity is a process that occurs postnatally in mammals as a continuation of β cell development, however an unified mechanistic model of β cell maturation is still missing [37]. In comparison to native islets, the stem cell-derived β cells do not undergo full functional maturation, which can be reflected by the glucose threshold responsiveness that triggers insulin secretion [38, 39]. Otherwise, several studies delineated the progression steps along the pancreatic differentiation path and identified the transcriptional signature determining endocrine cell fate and its early maturation sequence during ontogenesis [37, 40, 41]. In particular, time-specific onset of transcription factors like *NEUROD1*, *NKX6.1*, *NKX2.2* and *MNX1* enables the endocrine differentiation [42–44] as well as the differential expression of *ISL1* and *FEV* dictates the commitment of polyhormonal-fated cells towards INS^+^/PCSK1^+^ β-like cells, ARX^+^/GCG^+^ α-like cells or other hormone-producing cells [27, 43, 45].

With reference to the transcriptional identity of stem cell-derived endocrine cells and the kinetics of β cell maturity markers as they are codified in the literature, we found that WFS1 iBeta had a transcriptional fingerprint characterized by high levels of the early endocrine markers *NEUROD1* and *MAFB*, and by *ISL1* expression preceding *FEV* initiation that also marks polyhormonal-fated cells [27, 46]. Indeed, we found a significant enrichment of INS^+^/GCG^+^/SST^+^ or INS^+^/GCG^+^/PPY^+^ polyhormonal cells in WFS1 iBeta compared to its gene engineered counterpart. According to the relative percentage of polyhormonal cells, WFS1 iBeta showed lower levels of α-, PP- and δ-like cells. Conversely, they displayed increased mono-hormonal INS^+^ β-like cells, although the genetic correction to restore WT Wolframin led to iBeta with higher expression of *NKX6.1*, *NKX2.2* and *MNX1* endocrine markers. However, despite the increase of INS^+^ β-like cells in WFS1 iBeta, we showed that differentiation to β-like cells of patient-derived unedited iPSCs hesitated in a step backwards from a maturation and functional point of view, compared to gene edited cells, as supported by reduced expression of *UCN3*, *SYT4*, *CFPA126* and *MAFA* [40, 47–49]. In particular, *SYT4* expression increases during β cell maturation to modulate the Ca^2+^ sensitivity of secretory granules [50]. Analogously, *CFPA126* gene (also known as Flattop or Fltp) is involved in insulin secretion and glucose responsiveness [51], and its expression is correlated with WNT-signaling pathway and the expression of *WNT4* gene, which contributes to the paracrine signals between the mature β cells to control the activation of Ca^2+^ signaling in response to a glucose challenge [52]. Indeed, we found that the expression of WNT4 was significantly reduced in WFS1 iBeta. Together with lower expression of *SYT4*, *CFPA126* and *MAFA*, this data could explain the impairments in Ca^2+^ fluxes and glucose-stimulated insulin secretion (GSIS) observed in WS1-affected iBeta. However, in addition to an insufficient expression of genes involved in β cell maturation, we also found a significant decrease in expression of *SNAP25*, a key marker of the secretory machinery. In pancreatic β cells, the glucose-driven insulin secretion is mediated by the SNARE complex, which includes the plasma membrane proteins synaptotagmins (Syt7 and Syt4) and SNAP25, and the vesicular protein VAMP2/synaptobrevin [53, 54]. As the increase of Syt4 expression over Syt7 ameliorates glucose sensitivity and increases during β cell maturation [50], expression of *SNAP25* and *VAMP2* should occur at the same time during endocrine development. Since *VAMP2* expression did not change between WFS1 and WFS1^wt/757A>T^ iBeta, we hypothesized that *SNAP25* downmodulation might be independent of the maturation degree of WS1-affected iBeta and be correlated with Wolframin defects. Relative WFS1 β cell immaturity and reduced expression of *SYT4*, *WNT4* and *SNAP25* explained improper glucose responsiveness, delayed β cell inactivation upon stimulation and Ca^2+^ fluxes alterations. Specifically, the downregulation of *SNAP25* has been reported to correlate with desynchronization of Ca^2+^ oscillations [25]. Reduced SNAP25-dependent β cell co-activity and Syt4-related high Ca^2+^ influx sensitivity should hesitate in increasing insulin secretion, but we found that insulin release AUC was significantly lower in WFS1 iBeta in comparison to their gene edited counterpart. This result, however, is not contradictory, as we validated in WFS1 iBeta a reduced expression of *CACNA1D* gene, which encode the L-type Ca^2+^ channel Ca_v_1.3 that positively regulates GSIS [10], and an increase in the expression of *RGS4* gene, a negative regulator of insulin release [30]. Fewer Ca_v_1.3 channels and increased *RGS4* act as a compensatory mechanism that limits the amount of insulin secreted after glucose challenge. Furthermore, also the reduced amount of PC1/3 at both transcriptional and protein levels in the WS1-affected β cells, mainly correlating with their degree of maturity [55], could explain the lower insulin levels observed in dynamic perifusion studies. We also examined whether insulin processing differed between WFS1 and WFS1^wt/757A>T^ iBeta cells by measuring the ratio of secreted proinsulin to insulin. The increased proinsulin levels in WFS1 cells may result from the PC1/3 defect and ER stress [13–15].

Autophagy is an essential process to maintain the proper architecture, mass and function of pancreatic β cells, and its alterations are responsible for insulin deficiency [56]. Autophagy protects from apoptosis, preserves insulin secretory granules and maintains mitochondrial function, contributing to the homeostatic control of β cells [57]. Recently, “secretory autophagy” gained attention as unconventional mechanisms by which selected cargos are carried by the double-membrane autophagosome and exported to the extracellular environment for exocytosis [58]. Increased autophagic activity was reported to induce insulin secretion in β cells in a Ca^2+^-dependent manner [59]. In our case, the partially functional Wolframin with consequent Ca^2+^ dysfunction might impact the insulin secretion mediated by the autophagic vesicles. Moreover, the increase in autophagic flux we observed in WFS1 iBeta compared to WFS1^wt/757A>T^ counterpart might be a consequence of the accumulation of secretory granules that cannot be released from the cell due to the impairments we described above, thus undergoing degradation. However, further colocalization analyses between autophagic and lysosomal markers and insulin granules are required to confirm this hypothesis.

The clinical heterogeneity of WS1 poses challenges to the design of effective therapeutic strategies to treat the disorder. Drug-repurposing approaches have been exploited to target multiple mechanisms to ultimately alleviate the symptoms, without delaying or solving the pathology [17]. The class of GLP-1R agonists was shown to provide beneficial effects as disease-modifying agents in both *in vitro* and preclinical models, as well as in clinical settings. Indeed, the patient of the present study initiated an off-label liraglutide treatment that increased the levels of c-peptide, leading to the reduction of the total daily insulin dose, while the neuro-ophthalmology parameters remained stable [18].

According to results obtained in our clinical trial, we demonstrated that liraglutide effectively corrects insulin secretion ability and significantly increases GSIS when added together with glucose during dynamic perifusion. Despite previous works suggesting that anti-hyperglycemic effects of liraglutide are pleiotropically expected as extra-pancreatic action [60], more recent publication showed that the liraglutide-induced insulin secretion is related to the stimulation of the hyperpolarization-activated cyclic nucleotide-gated (HCN) channels on β and δ cells, that increase Ca^2+^ oscillation frequency in islets [61]. Interestingly, we extended this evidence, confirming that liraglutide treatment was able to resynchronize β cells by modulating Ca^2+^ fluxes. Moreover, liraglutide demonstrated its efficacy in restoring Ca^2+^ oscillations in TG-treated WFS1 and WFS1^wt/757A>T^ cells, supporting the hypothesis that GLP-1 signaling may play a role in Ca^2+^-dependent exocytosis of insulin granules. Additionally, the observed upregulation of *CACNA1D* and *SNAP25* further underscores the therapeutic long-lasting potential of liraglutide, as it would seem to reprogram the GSIS-related transcriptional landscape, positively influencing the molecular machinery involved in insulin secretion. Indeed, GLP-1 was previously shown to induce the phosphorylation of SNAP25 in β cells, with a putative role on downstream insulin secretion capacity [62]. The activation of the GLP-1 axis has been reported to promote DNA binding of PDX1 and the transcription of INS [63], and GLP-1 treatment is a potent inducer of β cell terminal differentiation [64]. Furthermore, liraglutide was found to be able to upregulate *MAFA*, *INS* and *PCSK1* genes, promoting a more “β-like state” [65]. Collectively, our findings support the growing body of evidence that GLP-1R agonists like liraglutide not only improve glycemic control through their insulinotropic effects, but also ameliorate β cell function at a cellular and molecular level.

Finally, chronic use of liraglutide exerted antiapoptotic effects, protecting the WFS1 iBeta from both TG and inflammation-induced apoptosis, without apparently alleviating ER stress [19] as we conversely observed a significant increase of UPR markers in the liraglutide-treated cells. This is in accordance with previous evidence where GLP-1R agonists displayed the capacity to reduce apoptosis in both β cells and neurons [19, 66, 67]. Specifically, we observed increased expression of chaperon proteins, such as BiP and PDIA4, at levels comparable to those observed in genetically corrected lines. The overexpression of these markers seemed to correlate with the increased and prolonged response to exogenous stress and the ability to compensate for the alterations deriving from the accumulation of aberrant transcripts and misfolded proteins that we observed in the cells deriving from the patient object of the present study. Restoring the ability to adapt and recover from chronic stress enforces intrinsic metabolic plasticity that β cells cyclically display to overcome ER stress and prevent apoptosis [68]. Interestingly, our insights are supported by investigation demonstrating anti-inflammatory properties of GLP-1R agonists [69]. Moreover, it has been reported that liraglutide mitigated neuroinflammation and reduced levels of circulating TNF-α, IL-1β, and IL-6, ameliorating lung injury and modulating immune cells in mouse models [70]. Here we demonstrated that the anti-inflammatory features of liraglutide also rely on the capacity to reduce susceptibility to inflammatory-mediated damage of β cells, paving the way to support the possibility of the therapeutic use of GLP-1R agonists in immune-mediated diabetes as well. As the patient of the present study showed high levels of systemic inflammation that we demonstrated may strongly affect survival of β cells, the use of GLP-1R agonists could represent the best therapeutic options since it was able to restore insulin secretion and protect β cell from chronic systemic inflammation resulting from these specific *WFS1* variants.

Our study highlights the relevance of unveiling the alterations of pathways associated with specific *WFS1* mutations in order to better elucidate the polyfunctional roles of Wolframin in pancreatic β cells. These insights are necessary to uncover molecular targets associated with specific *WFS1* variants to design more effective therapeutics according to a precision-medicine approach.

## Data availability

All raw data that were not directly included in the manuscript or that have not been deposited in online repositories, are available on request from the corresponding authors.

## Supporting information

ESM Figure 1

ESM Figure 2

ESM Figure 3

ESM Figure 4

ESM Tables

## Acknowledgments

We thank the Advanced Light and Electron Microscopy BioImaging Center (ALEMBIC), at San Raffaele Scientific Institute, Milan (Italy), for confocal immunofluorescence and calcium flux assay images, and the Flow cytometry Resource, Advanced Cytometry Technical Applications Laboratory (FRACTAL), at San Raffaele Scientific Institute, Milan (Italy), for FACS analysis. We also thank Dr. Francesca Giannese and Dr. Dejan Lazarevic at Center for Omics Sciences (COSR) of the San Raffaele Scientific Institute, Milan (Italy), for having provided support in library preparation and scRNAseq experiments, and Dr. Giulia Scotti for scRNAseq sample quality control and preliminary bioinformatic analysis. Dr. Silvia Torchio conducted this study as partial fulfillment of an international PhD in Molecular Medicine at Vita-Salute San Raffaele University.

## Author contributions

Conceptualization and experimental design, R.C. and L.P.; methodology, S.T. and R.C.; investigation, S.T., G.S., F.C., V.Z., S.P., F.M., R.C.; bioinformatic analysis, R.C.; formal analysis and interpretation of results, S.T., G.S., R.C., and L.P.; stem cell differentiation, F.C., V.Z., S.P., and V.S.; writing - original draft, S.T., G.S. and R.C.; writing - review and editing, S.T., G.S., R.B., G.F., V.S., R.C. and L.P.; supervision, R.C. and L.P.; responsible for the integrity of the work as a whole, L.P.

**ESM Figure 1 (a)** The count/barcode curve for WFS1 iBeta. Blue line indicates the number of barcodes estimated by the tools (n=4594). **(b)** Distribution of the number of genes, number of UMIs and percentage of mitochondrial genes/cell in WFS1 iBeta. **(c)** UMAP scoreplot for WFS1 iBeta, color-coded according to the cell cycle phases (G1=pink; G2M=green; S=blue) and to the number of expressed genes/cell (purple blue gradient). **(d)** The count/barcode curve for WFS1^wt/757A>T^ iBeta. Blue line indicates the number of barcodes estimated by the tools (n=2215). Green line indicates the number of barcodes arbitrarily extracted (n=3000). **(e)** Distribution of number of genes, number of UMIs and percentage of mitochondrial genes/cell in WFS1^wt/757A>T^ iBeta. **(f)** UMAP scoreplot for WFS1^wt/757A>T^ iBeta, color-coded according to the cell cycle phases (G1=pink; G2M=green; S=blue) and to the number of expressed genes/cell (purple blue gradient). Heatmap representing the expression of the top 10 marker genes (according to log fold change) identified for each cluster (0-6) among all the cells of **(g)** WFS1 and **(h)** WFS1^wt/757A>T^ iBeta.

**ESM Figure 2** Percentages of mono-hormonal (α, β, δ and PP) and poly-hormonal cells in the endocrine clusters of **(a)** WFS1 and **(b)** WFS1^wt/757A>T^ iBeta. **(c)** Violin plot detailing log-normalized gene expression of late β cell maturation markers *UCN3*, *SYT4*, *CFAP126* and *MAFA* in WFS1 (red) and WFS1^wt/757A>T^ (green) endocrine clusters. *p < 0.05, ***p < 0.001 by Mann-Whitney nonparametric test. **(d)** Pivotal transcription factors (TFs) regulating both β cell differentiation and maturation, following schematic representation of their expression onset and timing. Color gradients indicate the increase/decrease in the expression levels over time.

**ESM Figure 3** Relative expression of *WFS1* gene in both basal conditions (DMSO) and after treatment with either 50 or 500 nM TG in WFS1 (red) and WFS1^wt/757A>T^ (green) iBeta. Data are plotted as mean±SD and expressed as fold over DMSO-treated cells. N=5. *p<0.05 by two-way ANOVA with Dunn-Šídák correction.

**ESM Figure 4 (a)** Mean fluorescence signals (F/F_0_) over time indicating [Ca^2+^]_i_ oscillations in WFS1 β cells treated with 50 nM TG following 11 mM glucose stimulation without (red) or with (blue) 1 μM liraglutide. The average and maximum peaks values are reported. N=20 from 2 independent experiments. **(b)** Mean fluorescence signals (F/F_0_) over time indicating [Ca^2+^]_i_ oscillations in WFS1^wt/757A>T^ β cells treated with 50 nM TG following 11 mM glucose stimulation without (green) or with (blue) 1 μM liraglutide. The average and maximum peaks values are reported. N=20 from 2 independent experiments.

**ESM Table 1** List of antibodies employed in the study and respective mode of use.

**ESM Table 2** List of the 48 genes analyzed with the TaqMan Array microfluidic card.

**ESM Table 3** List of primers used for 96-well RT-qPCR experiments.

