## Supplementary figures and images for "Liraglutide treatment reverses unconventional cellular defects in induced pluripotent stem cell-derived β cells harboring a partially functional WFS1 variant"

### ESM Figure 1

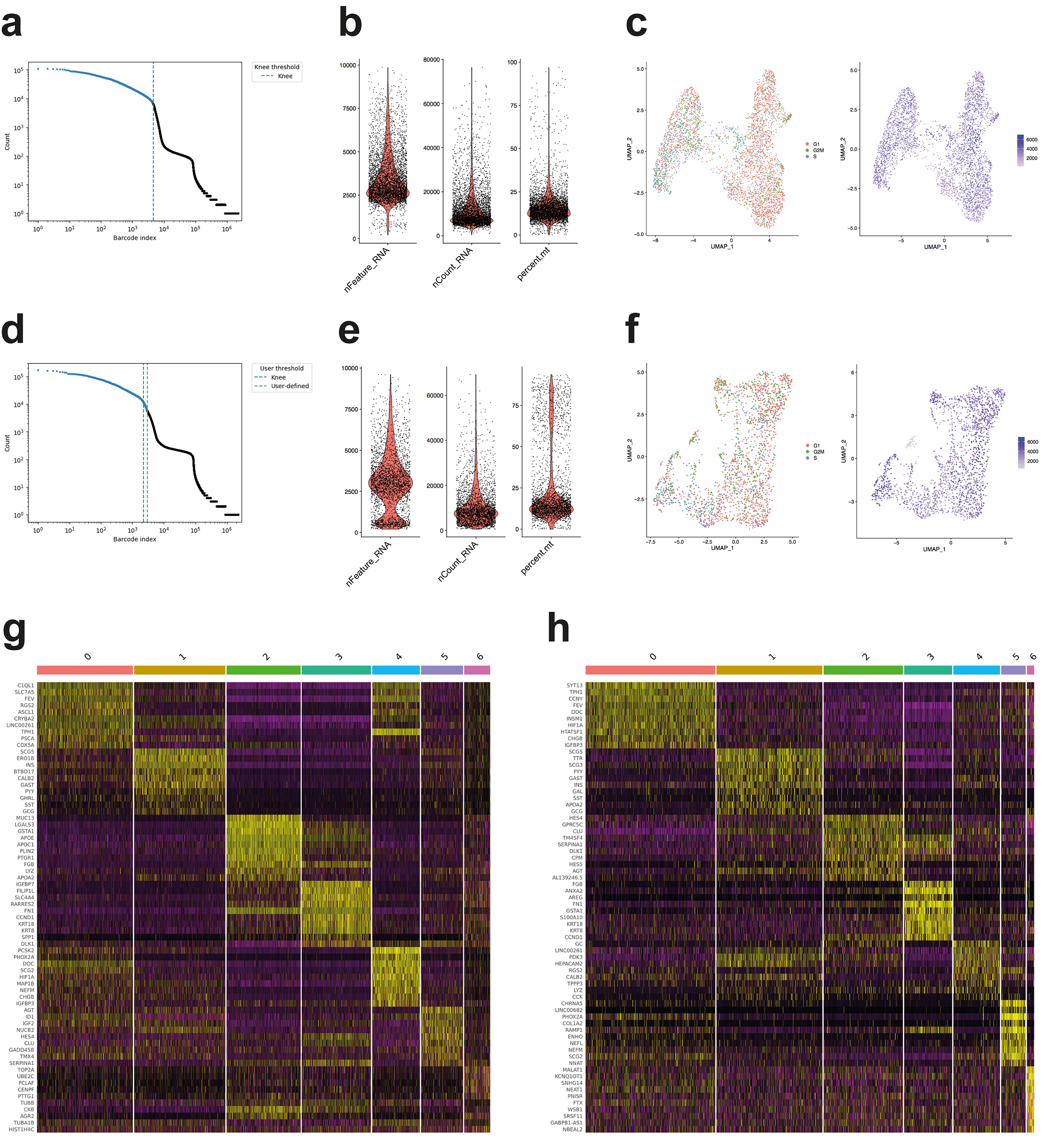

### ESM Figure 2

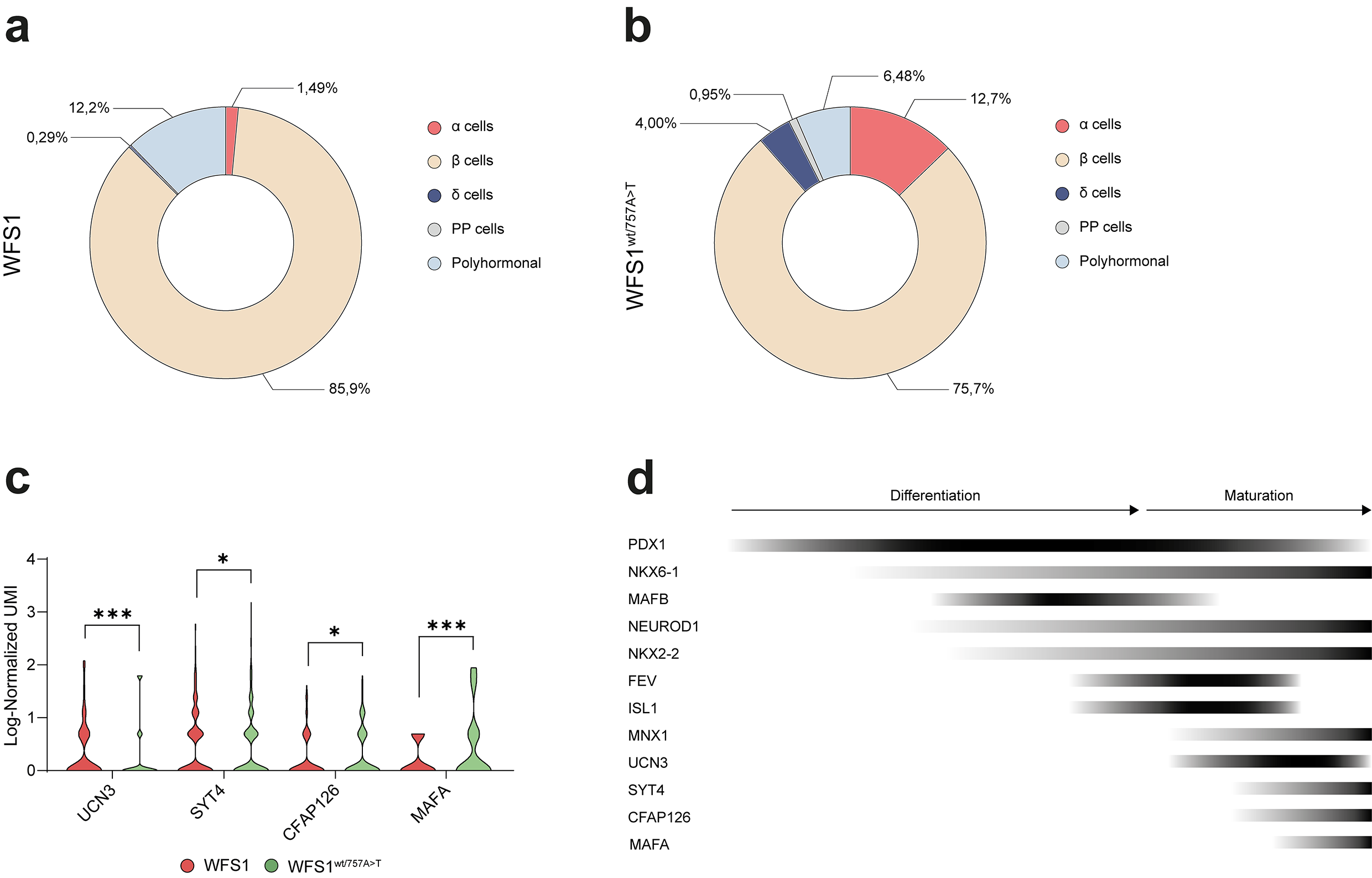

### ESM Figure 3

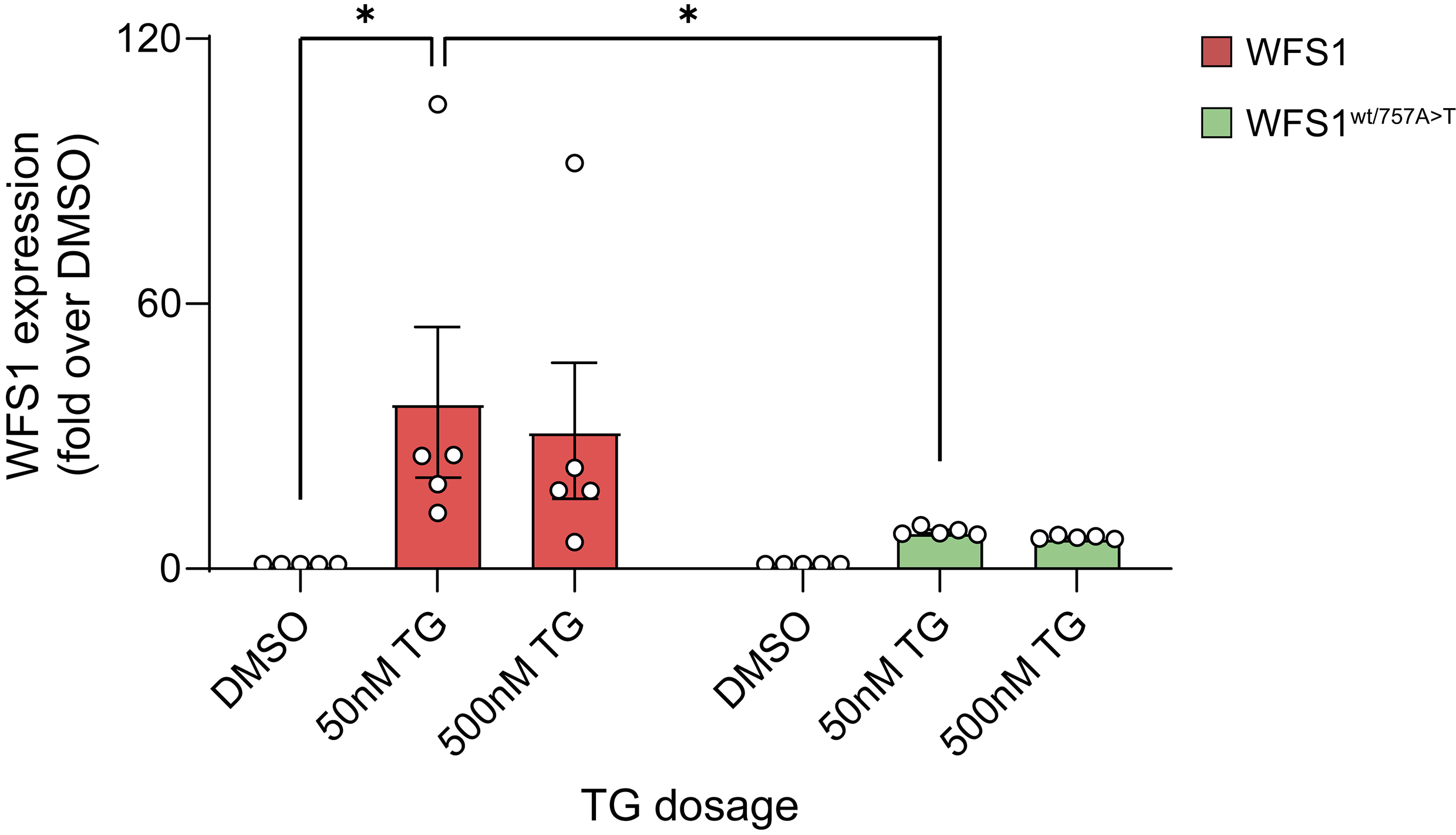

### ESM Figure 4

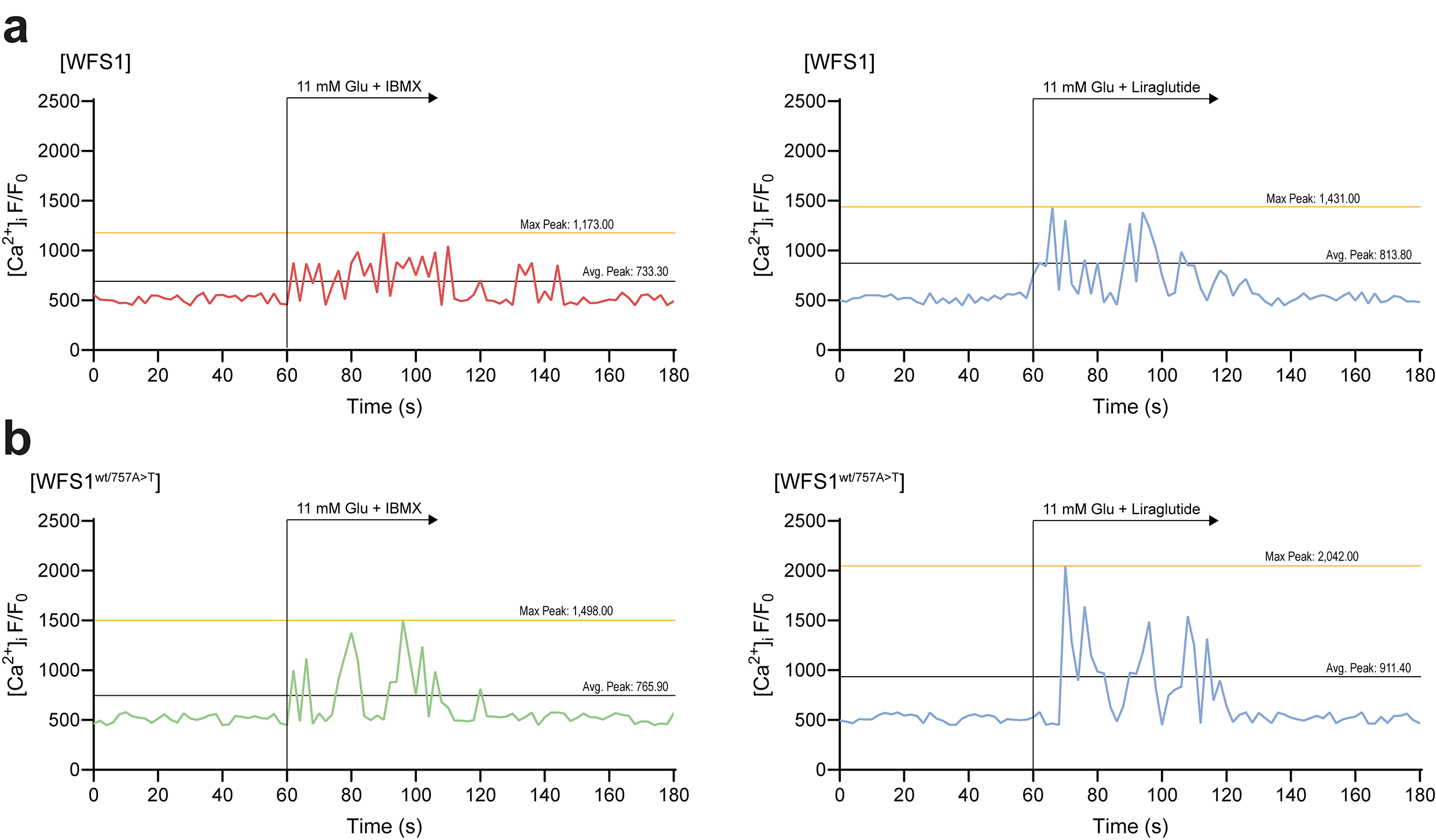
