## Supplementary material for "Liraglutide treatment reverses unconventional cellular defects in induced pluripotent stem cell-derived β cells harboring a partially functional WFS1 variant": ESM Tables

**ESM Table 1. List of antibodies employed in this study and respective mode of use.**

| **Antibody** | **Species** | **Supplier** | **Code** | **Use** | **Dilution** |
| --- | --- | --- | --- | --- | --- |
| GAPDH [6C5] | Mouse | Abcam | ab8245 | WB | 1:5000 |
| Alexa Fluor® 647  anti-Insulin [T56-706] | Mouse | BD | T56-706 Alexa-Fluor647 | FACS | 2µl x 10^6^ cells |
| Insulin | Guinea pig | Dako | A0564 | IF | 1:200 |
| LC3A/B [D3U4C] | Rabbit | Cell Signaling | #12741 | WB | 1:1000 |
| PE anti-NKX6.1 [R11-560] | Mouse | BD | R11-560 PE | FACS | 5µl x 10^6^ cells |
| NKX6.1 [631438] | Mouse | R&D | MAB5857 | IF | 1:100 |
| PC1/3 [EPR21908] | Rabbit | Abcam | ab220363 | WB | 1:1000 |
| Alexa Fluor® 488 anti-PDX1 [658A5] | Mouse | BD | 658A5 Alexa-Fluor488 | FACS | 5µl x 10^6^ cells |
| SQSTM1 [2C11] | Mouse | Abcam | ab56416 | WB | 1:1000 |
| HRP-conjugated anti-mouse | Goat | R&D | HAF007 | WB | 1:1000 |
| HRP-conjugated anti-rabbit | Goat | R&D | HAF008 | WB | 1:1000 |
| Alexa Fluor® 647  anti-Guinea Pig | Goat | ThermoFisher | A-21450 | IF | 1:500 |
| Alexa Fluor® 546 anti-Mouse | Goat | ThermoFisher | A-11030 | IF | 1:500 |

**ESM Table 2. List of the 48 genes included in the TaqMan Array microfluidic card.**

| GAPDH-Hs99999905_m1 | glyceraldehyde-3-phosphate dehydrogenase |
| --- | --- |
| INS-Hs02741908_m1 | insulin |
| CHGA-Hs00900375_m1 | chromogranin A (parathyroid secretory protein 1) |
| SYP-Hs00300531_m1 | synaptophysin |
| PTF1A-Hs00603586_g1 | pancreas specific transcription factor, 1a |
| NEUROG3-Hs01875204_s1 | neurogenin 3 |
| NEUROD1-Hs00159598_m1 | neuronal differentiation 1 |
| POU5F1-Hs04260367_gH | POU class 5 homeobox 1 |
| PAX4-Hs00173014_m1 | paired box 4 |
| ONECUT1-Hs00413554_m1 | one cut homeobox 1 |
| ISL1-Hs00158126_m1 | ISL LIM homeobox 1 |
| GAD2-Hs00609534_m1 | glutamate decarboxylase 2 (pancreatic islets and brain, 65kDa) |
| PAX6-Hs00240871_m1 | paired box 6 |
| SOX17-Hs00751752_s1 | SRY (sex determining region Y)-box 17 |
| GCG-Hs01031536_m1 | glucagon |
| PDX1-Hs00236830_m1 | pancreatic and duodenal homeobox 1 |
| NKX6-1-Hs00232355_m1 | NK6 homeobox 1 |
| NKX2-2-Hs00159616_m1 | NK2 homeobox 2 |
| NANOG-Hs04399610_g1 | Nanog homeobox |
| MAFA-Hs04419862_g1 | v-maf avian musculoaponeurotic fibrosarcoma oncogene homolog A |
| FOXA2-Hs00232764_m1 | forkhead box A2 |
| HNF1B-Hs00172123_m1 | HNF1 homeobox B |
| SOX2-Hs01053049_s1 | SRY (sex determining region Y)-box 2 |
| ARX-Hs00292465_m1 | aristaless related homeobox |
| FOXO1-Hs01054576_m1 | forkhead box O1 |
| RFX6-Hs00543100_m1 | regulatory factor X, 6 |
| FGF10-Hs00610298_m1 | fibroblast growth factor 10 |
| NOTCH1-Hs01062014_m1 | notch 1 |
| HHEX-Hs00242160_m1 | hematopoietically expressed homeobox |
| SLC2A2-Hs01096900_g1 | solute carrier family 2 (facilitated glucose transporter), member 2 |
| ONECUT3-Hs01369570_m1 | one cut homeobox 3 |
| FOXA3-Hs00270130_m1 | forkhead box A3 |
| MNX1-Hs00907365_m1 | motor neuron and pancreas homeobox 1 |
| INSM1-Hs00357871_s1 | insulinoma-associated 1 |
| PCSK1-Hs01026107_m1 | proprotein convertase subtilisin/kexin type 1 |
| SNTB2-Hs00198844_m1 | syntrophin, beta 2 |
| SST-Hs00356144_m1 | somatostatin |
| GHRL-Hs01074053_m1 | ghrelin/obestatin prepropeptide |
| PPY-Hs00358111_g1 | pancreatic polypeptide |
| GCK-Hs01564555_m1 | glucokinase (hexokinase 4) |
| IAPP-Hs00169095_m1 | islet amyloid polypeptide |
| ABCC8-Hs01093761_m1 | ATP-binding cassette, sub-family C (CFTR/MRP), member 8 |
| SLC30A8-Hs00545183_m1 | solute carrier family 30 (zinc transporter), member 8 |
| PTPRN-Hs01090891_g1 | protein tyrosine phosphatase, receptor type, N |
| TSPAN7-Hs00190284_m1 | tetraspanin 7 |
| TMEM27-Hs00252907_m1 | transmembrane protein 27 |
| G6PC2-Hs01549773_m1 | glucose-6-phosphatase, catalytic, 2 |
| KCNJ8-Hs00958961_m1 | potassium inwardly-rectifying channel, subfamily J, member 8 |

**ESM Table 3. List of PCR primers employed in this study.**

| ***Gene* (Protein)** | **Forward** | **Reverse** |
| --- | --- | --- |
| *ATF4* | GTTCTCCAGCGACAAGGCTA | ATCCTGCTTGCTGTTGTTGG |
| *ATG10* | GGTGATAGTTGGGAATGGAGACC | GTCTGTCCATGGGTAGATGCTC |
| *ATG12* | GGGAAGGACTTACGGATGTCTC | AGGAGTGTCTCCCACAGCCTTT |
| *BECN1* | CTGGACACTCAGCTCAACGTCA | CTCTAGTGCCAGCTCCTTTAGC |
| *CACNA1C* | GGACGTGCTGTACTGGATGC | GGCCTTCTCCCTCTCTTTGG |
| *CACNA1D* | CTTCGACAACGTCCTCTCTGCT | GCCGATGTTCTCTCCATTCGAG |
| *DDIT3* (CHOP) | AGAACCAGGAAACGGAAACAGA | TCTCCTTCATGCGCTGCTTT |
| *GAPDH* | GTCTCCTCTGACTTCAACAGCG | ACCACCCTGTTGCTGTAGCCAA |
| *HERPUD1* | TACTCCTCCCTGAGCAGATTCC | TTTCAGGATCAGTGCCTTCCTGT |
| *HSPA5* (BiP) | TGTTCAACCAATTATCAGCAAACTC | TTCTGCTGTATCCTCTTCACCAGT |
| *PCSK1* | GCTGGGCTATGACCTTTTGG | CAGCCCATATCACACGATCA |
| *PDIA4* | CTCCAGAACCCAGGAAGAAATTG | CTTCTCATACTCGGGGGCAA |
| *SEL1L* | GTGGGGCTTTTGTGAAACTGAA | TGACACTCTCTCCAGGGCTT |
| *SNAP25* | CGTCGTATGCTGCAACTGGTTG | GGTTCATGCCTTCTTCGACACG |
| *SQSTM1* | TGCCCAGACTACGACTTGTG | AGTGTCCGTGTTTCACCTTCC |
| *VAMP2* | CTCCAAACCTCACCAGTAACAGG | AGCTCCGACAGCTTCTGGTCTC |
| *WNT4* | GAGCAACTGGCTGTACCTGG | TGAGTTTCTCGCACGTCTCC |
| *WFS1* | GGGCCTACAAAGGGAGACAT | CCAGTACATGACCAGGGCTG |
